# Two sides of a coin: a Zika virus mutation selected in pregnant rhesus macaques promotes fetal infection in mice but at a cost of reduced fitness in nonpregnant macaques and diminished transmissibility by vectors

**DOI:** 10.1101/2020.08.11.247411

**Authors:** Danilo Lemos, Jackson B. Stuart, William Louie, Anil Singapuri, Ana L. Ramírez, Jennifer Watanabe, Jodie Usachenko, Rebekah I. Keesler, Claudia Sanchez-San Martin, Tony Li, Calla Martyn, Glenn Oliveira, Sharada Saraf, Nathan D. Grubaugh, Kristian G. Andersen, James Thissen, Jonathan Allen, Monica Borucki, Konstantin A. Tsetsarkin, Alexander G. Pletnev, Charles Y. Chiu, Koen K. A. Van Rompay, Lark L. Coffey

## Abstract

Although fetal death is now understood to be a severe outcome of congenital Zika syndrome, the role of viral genetics is still unclear. We sequenced Zika virus (ZIKV) from a rhesus macaque fetus that died after inoculation and identified a single intra-host mutation, M1404I, in the ZIKV polyprotein, located in NS2B. Targeted sequencing flanking position 1404 in 9 additional macaque mothers and their fetuses identified M1404I at sub-consensus frequency in the majority (5 of 9, 56%) of animals and some of their fetuses. Despite its repeated presence in pregnant macaques, M1404I occurs rarely in humans since 2015. Since the primary ZIKV transmission cycle is human-mosquito-human, mutations in one host must be retained in the alternate host to be perpetuated. We hypothesized that ZIKV I1404 increases fitness in non-pregnant macaques and pregnant mice but is less efficiently transmitted by vectors, explaining its low frequency in humans during outbreaks. By examining competitive fitness relative to M1404, we observed that I1404 produced lower viremias in non-pregnant macaques and was a weaker competitor in tissues. In pregnant wildtype mice ZIKV I1404 increased the magnitude and rate of placental infection and conferred fetal infection, contrasting with M1404, which was not detected in fetuses. Although infection and dissemination rates were not different, *Ae. aegypti* transmitted ZIKV I1404 more poorly than M1404. Our data highlight the complexity of arbovirus mutation-fitness dynamics, and suggest that intrahost ZIKV mutations capable of augmenting fitness in pregnant vertebrates may not necessarily spread efficiently via mosquitoes during epidemics.

**IMPORTANCE:** Although Zika virus infection of pregnant women can result in congenital Zika syndrome, the factors that cause the syndrome in some but not all infected mothers are still unclear. We identified a mutation that was present in some ZIKV genomes in experimentally inoculated pregnant rhesus macaques and their fetuses. Although we did not find an association between the presence of the mutation and fetal death, we performed additional studies with it in non-pregnant macaques, pregnant mice, and mosquitoes. We observed that the mutation increased the ability of the virus to infect mouse fetuses but decreased its capacity to produce high levels of virus in the blood of non-pregnant macaques and to be transmitted by mosquitoes. This study shows that mutations in mosquito-borne viruses like ZIKV that increase fitness in pregnant vertebrates may not spread in outbreaks when they compromise transmission via mosquitoes and fitness in non-pregnant hosts.

## INTRODUCTION

Congenital Zika syndrome (CZS) caused by Zika virus (ZIKV) produces a disease spectrum that sometimes results in microcephaly or death in fetuses from mothers infected during pregnancy. In American outbreaks since 2015, about 15% of fetuses from ZIKV infected mothers displayed reduced growth, sensory disorders, and central nervous system malformations (1–6), manifestations of CZS (7–9). CZS abnormalities associate with detection of ZIKV RNA or infectious virus in amniotic fluid (AF) and fetal tissues, including brain (4–6, 10– 14). Viral and host factors that affect the severity of CZS are still not well understood, including why some fetuses die or develop microcephaly while others do not.

Mutations in ZIKV that arise and spread in humans during outbreaks may contribute to CZS or modify transmission by mosquitoes. However, identifying mutations that could influence ZIKV phenotype is complicated as consensus (average nucleotide) genomes from febrile human cases in recent outbreaks differ by hundreds of nonsynonymous mutations compared to ancestral genomes (15). Any of these mutations alone or in combination could modify incidence, transmissibility, or pathogenesis. Furthermore, the consensus genomes from 1 miscarriage and 7 microcephaly cases are interleaved with febrile genomes in phylogenies and share no common amino acid differences compared to febrile cases from patients living in the same areas suggesting that no single mutation or group of mutations associate with the most severe CZS outcomes (16). Phylogenetic inference identified prM protein (prM-S139N) coding mutation that increases microcephaly in mice (17), although this finding was not reproducible in repeated studies (18). Another mutation identified via phylogenetic analysis, E-V473M, increased viremias in non-pregnant cynomolgus macaques and neurovirulence in 1-day old mice inoculated intracranially, but did modify ZIKV RNA levels in *Ae. aegypti* (19). Since prM-S139N and E-V473M are present in most genomes from American outbreaks since 2015, including in febrile women whose babies did not develop CZS, they are likely not major determinants of CZS outcome in pregnant women.

ZIKV evolution intrahost has been less studied than evolution across patients but may also affect disease outcome. Defining intrahost evolution over time necessitates repeated sampling. For pregnant women, sequencing ZIKV from even a single time is often unsuccessful, since viremia is frequently missed in the clinical setting (14, 20), and AF is rarely available given that amniocentesis can lead to iatrogenic infection (21) or ZIKV transmission to the fetus from an infected mother. Due to these limitations, the role of intrahost viral genetics in CZS remains unclear. To circumvent limited human sample availability or testing hundreds of mutations identified via phylogenetics as potential determinants of CZS, we used an experimental model to identify intrahost ZIKV mutations in infected pregnant rhesus macaques with known CZS outcomes.

Macaques have rapidly become an important model for understanding ZIKV infection and disease (22, 23, 32–34, 24–31) due to similar placentation, immunology, fetal organogenesis, and neurologic development with humans. Studies using rhesus macaques, including our own work (24, 35), have demonstrated fetal central nervous system lesions consistent with abnormal brain development observed in CZS. As in humans, not all rhesus macaque fetuses from mothers inoculated with the same ZIKV stock at similar gestational windows develop CZS. While host determinants certainly play a role, viral mutations developed intrahost may also influence different CZS outcomes. Sub-consensus mutants in genetically heterogeneous populations of many other RNA viruses (36, 37, 46–51, 38–45) have been associated with modified transmissibility or disease outcomes. A wild type 2015 ZIKV strain from a febrile patient in Brazil showed augmented replicative success in human placental and neural cells compared to its genetically homogenous infectious clone derivative (16). This highlights that ZIKV mutants intrahost play a role in infection kinetics, even in cell monocultures. However, sub-consensus ZIKV mutations in an individual host are unrecognized without deep sequencing that characterizes the entire population of viral RNA genomes instead of solely the consensus. Intrahost ZIKV populations from 11 pregnant rhesus macaques revealed no *de novo* mutations in one study (51), although sequencing from maternal serum and AF was from just one time point post-inoculation.

In this study, we performed intrahost ZIKV sequencing in a rhesus macaque mother and her deceased fetus, and identified a single sub-consensus mutation, M1404I in the NS2B coding region. We then focused sequencing at position 1404 in 9 additional pregnant rhesus macaques and their fetuses where we found that it was present at a minority frequency in 5 additional animals but not detectable or at ∼1% frequency in the inoculum. Given its rise in frequency, we hypothesized that the mutant confers increased fitness in rhesus macaques and, by extension, other vertebrates. We therefore performed parallel infections of M1404 and I1404 in non-pregnant rhesus macaques and pregnant mice as well as mixed infections of non-pregnant rhesus macaques. Since ZIKV is primarily mosquito-borne, viral genomes that evolve in vertebrates must be maintained in mosquitoes to persist in alternating vertebrate-mosquito-vertebrate cycling. To test this concept and to better understand why I1404 was detected at low frequency in humans during outbreaks despite increasing in frequency intrahost in pregnant rhesus macaques, we compared transmissibility of M1404 and I1404 by parallel infections of *Aedes aegypti* vectors.

## RESULTS

### A ZIKV mutation arises *de novo* or increases in frequency in experimentally inoculated pregnant rhesus macaques

Pregnant rhesus macaques were ZIKV-inoculated to study viral infection dynamics and CZS. Outcomes from those experiments are detailed in Coffey *et al.* and Van Rompay *et al*. (24, 34). The rhesus macaques in those studies were inoculated in their first or second trimesters of pregnancy intravenously (IV) and intraamniotically (IA) with 1×10^5^ plaque forming units (PFU) of ZIKV WT (SPH2015, a Brazilian strain, KU321639) or subcutaneously (SC) with 1×10^3^ PFU of ZIKV Puerto Rico 2015 (KU501215) (**Figure 1**). Some fetuses died pre-term while most survived to the end of the study, which was ∼10 days pre-term. To investigate intrahost dynamics we analyzed samples archived from those rhesus macaques by deep sequencing. We first sequenced the complete ZIKV genome in amniotic fluid (AF) from rhesus macaque #5388 whose fetus died 7 days post inoculation (dpi) and exhibited high ZIKV RNA levels in multiple maternal and fetal tissues [23]. We identified no consensus (found in >50% of RNA reads) mutations and only one sub-consensus (found in <50% of RNA reads) mutation, G4315A, which occurred at 18% frequency. The G4315A mutation results in a non-synonymous methionine (M) to isoleucine (I) substitution at amino acid position 1404 of the ZIKV polyprotein (with reference to WT strain SPH2015, KU321639) and is located in the non-structural protein 2B (NS2B), a co-factor for the flavivirus protease, NS3. Sequencing of the inoculum by 2 different laboratories flanking the G4315A variant locus at 5,705 and 2,296-fold revealed the mutant in 0.2% or 1.3% of the ZIKV RNA reads, respectively (**Supplemental Table 2**), at or below the reported error rates for short-read Illumina sequencing (52). We performed additional sequencing to determine whether M1404I was also present in other maternal or fetal tissues from rhesus macaque #5338 and also in 9 additional pregnant ZIKV-inoculated rhesus macaques with different CZS outcomes using a targeted sequencing approach flanking 1404. In addition to the AF where it was first detected, ZIKV M1404I was detected in the gestational sac, amniotic membrane, placenta, and vagina of macaque #5388. ZIKV M1404I was also detected at minority frequency in 4 additional pregnant rhesus macaques inoculated IV and IA with the WT Brazilian strain in multiple tissues including amniotic fluid, gestational sac, amniotic membrane, placenta, maternal vagina, fetal seminal vesicle, and maternal urine. M1404I was also detected in the spleen of one mother (#5731) inoculated SC with the isolate from Puerto Rico. The Puerto Rican ZIKV inoculum lacked the I1404 variant in any RNA reads sequenced at a coverage depth of 2,242-fold. The I1404 mutation was not detected in tissues from 4 additional pregnant rhesus macaques in the same studies (**Supplemental Table 2**). Although the mutation was detected in 2 rhesus macaques whose fetuses died, it was also found in 4 rhesus macaques whose fetuses survived to near-term, indicating it was not associated with fetal death. The *de novo* development of ZIKV M1404I and its increase in frequency from near-absence in the inoculum to presence within multiple tissues intrahost in six pregnant rhesus macaques inoculated by two routes with two different ZIKV strains suggested that M1404I may be a rhesus macaque adaptive mutation. To study the mutation in isolation, we generated infectious clone derived viruses that vary only at 1404 for comparative fitness experiments in cells, non-pregnant rhesus macaques, pregnant mice, and mosquitoes.

**Figure 1.**
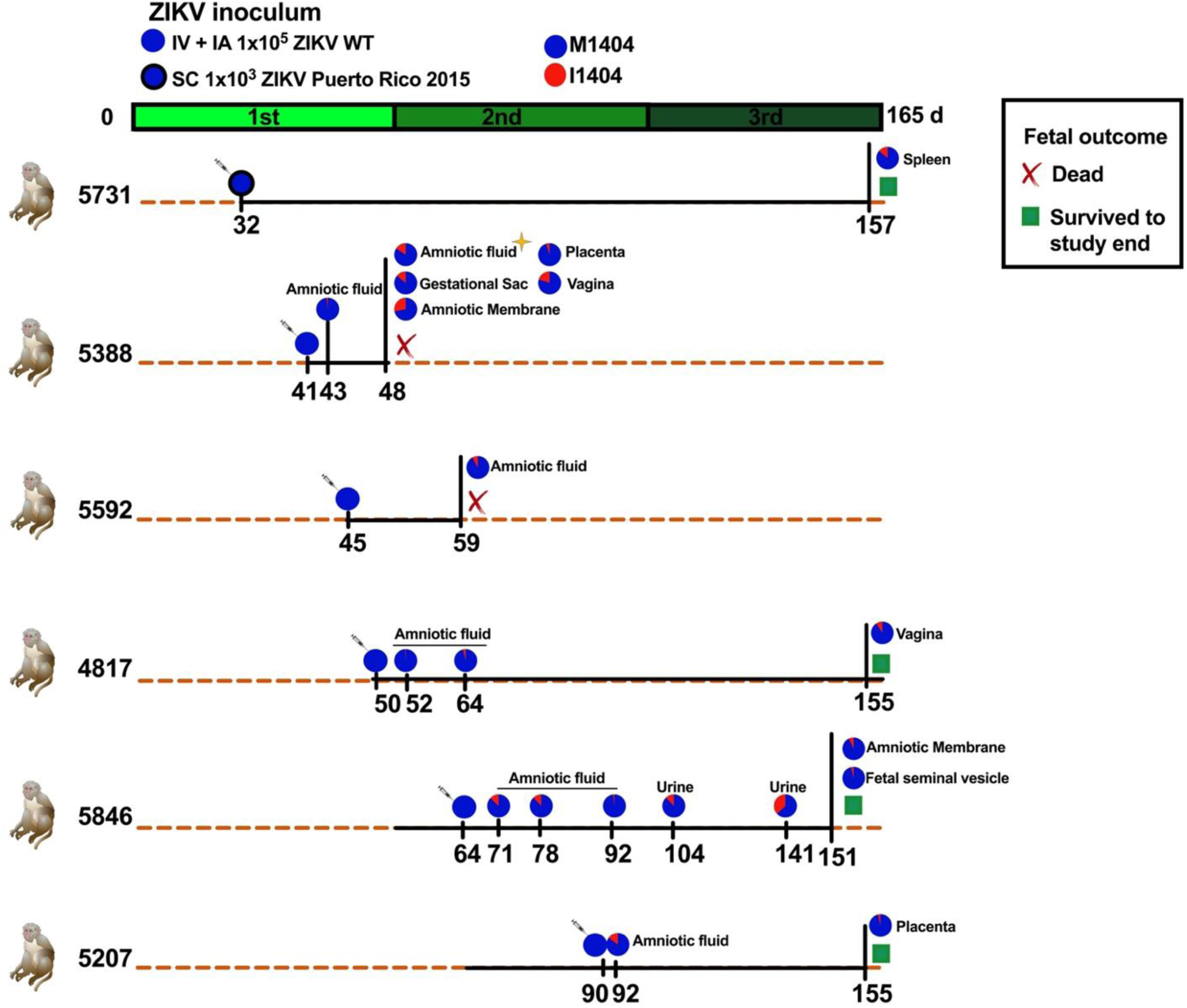
A ZIKV mutant arose or increased in frequency in six pregnant rhesus macaques inoculated with two different ZIKV strains via two routes. Experimental design for inoculation of pregnant rhesus macaques with 1×10^5^ ZIKV WT (KU321639.1, a 2015 Brazilian strain) both intravenously (IV) and intraamniotically (IA), blue circle, or 1×10^3^ of ZIKV Puerto Rico 2015 (KU501215.1) delivered subcutaneously (SC), blue circle with thick black border, on days indicated by syringes. For more details on study design, see Coffey *et al*. [23] and Van Rompay *et al.* [33]. Both inocula contained mostly M1404 where I1404 was absent or present at the limit of detection, ∼1%. The green line shows the duration of each rhesus macaque pregnancy divided into 3 ∼55 day trimesters where full-term is 165 days. The orange dotted lines represent 165 days of gestation and the black solid lines show the period of infection and experiment end for each dam and their fetus. Fetal death is shown as a red X. Fetuses that survived to the study endpoint, gestation day 155, are indicated with green squares. The golden star represents the first detection of I1404; full genome sequencing of this specimen showed no other genome-wide mutations. The pie charts represent the relative abundance of the amino acid at position 1404. The I1404 mutation was not detected in 4 additional pregnant macaques in the same study (not shown here but included in **Supplemental Table 2**). Supplemental Table 2 shows the depth of sequencing coverage for the data represented in pie charts. Tissues listed are maternal unless otherwise indicated.

### Growth kinetics of ZIKV I1404 are superior to M1404 in vertebrate cells

We tested whether expression of ZIKV I1404 in cell culture modifies ZIKV growth kinetics relative to M1404. We generated two infectious clones identical in sequence except for position 1404. To do this, we modified a ZIKV infectious clone made from a 2015 Brazilian ZIKV isolate, Paraiba_01/2015 (16), to match the amino acid sequence of the Brazilian strain SPH2015. Strain SPH2015, hereafter termed ‘WT’, was isolated from a patient and passaged in cells before use in pregnant rhesus macaque studies. The sequences of clone-derived ZIKV, termed ‘M1404’ and ‘I1404’ based on the amino acid at polyprotein position 1404 were verified by Sanger sequencing (data not shown) using primers flanking the ZIKV genome (**Supplemental Table 1)**. The relative growth kinetics measured as ZIKV RNA and infectious virus levels in supernatants of inoculated cultures over time were assessed in African green monkey kidney (Vero) or *Ae. albopictus* larval (C6/36) cells (**Supplemental Figure 1**). ZIKV I1404 exhibited significantly higher ZIKV RNA levels compared to M1404 from 24 to 96 hours post inoculation (hpi) in Vero cells at a MOI of 0.01 (p<0.001, repeated measures ANOVA) (**Supplemental Figure 1A**). ZIKV RNA levels for both M1404 and I1404 were lower than for WT at all times studied, a pattern common to clone-derived viruses compared to WT progenitor viruses [16]. Even though the increase in ZIKV RNA kinetics was accelerated for I1404 compared to M1404, infectious ZIKV levels in Vero cells from 24-96 hpi by plaque assay were not different between ZIKV M1404, I1404, and WT (p>0.05, repeated measures ANOVA) (**Supplemental Figure 1B**). In C6/36 mosquito cells, ZIKV RNA levels after inoculation at a MOI of 0.5 were not different across groups at any time from 24-96 hpi (p>0.05, repeated measures ANOVA) (**Supplemental Figure 1C**). These results indicate that in a standard vertebrate cell line, the I1404 mutation detected in pregnant rhesus macaques increases ZIKV RNA levels and infection kinetics compared to the progenitor M1404, although it does not change the levels of infectious virus.

### ZIKV I1404 displays reduced fitness in non-pregnant rhesus macaques inoculated with an equal mixture of M1404 and I1404

To directly compare the relative fitness of M1404 versus I1404, we performed a competition experiment wherein both viruses were inoculated together at equal ratios into two non-pregnant rhesus macaques (**Figure 2A**). Rhesus macaques were also inoculated with ZIKV WT or ZIKV I1404 alone for comparison. ZIKV M1404 was not included since its coding amino acid sequence is identical to WT to reduce macaque use. We defined fitness as the plasma viremia magnitude and kinetics and, for mixed infections, the relative abundance of ZIKV RNA reads encoding M1404 or I1404 in plasma and tissues. Rhesus macaques inoculated with ZIKV WT or the 1:1 mixture of M1404 and I1404 exhibited similar kinetics (p=0.474, unpaired t-test) with peak viremias at 5-7 dpi that reached 10^6^-10^7^ ZIKV genomes/ml plasma (**Figure 2B**) with no differences in the viremia area under the curve (AUC) (**Figure 2C**). By contrast, rhesus macaques inoculated with ZIKV I1404 developed significantly lower peak viremias, 10^2^ ZIKV genomes/mL plasma, that endured for a significantly shorter period of time and showed lower AUC (p=0.03, AUC) compared to the other two groups. Despite reduced magnitude and kinetics of viremia, most tissue ZIKV levels were not different in animals infected with I1404, WT, or the 1:1 mixture (**Figure 2D**). The I1404 mutant did not revert to M1404 in the spleen or mesenteric lymph node of either I1404-inoculated animal at levels detectable by Sanger sequencing (data not shown). Targeted deep sequencing flanking the 1404 locus in rhesus macaques inoculated with the 1:1 mixture detected ZIKV I1404 at a significantly lower frequency than M1404 in plasma, spleen, ileum, mesenteric, and inguinal lymph nodes (p<0.001, Chi-squared tests) (**Figure 2E, Supplemental Table 3**). Despite the repeated detection of ZIKV I1404 in pregnant rhesus macaques, these experiments show that I1404 reduces viral fitness relative to M1404 in non-pregnant rhesus macaques. Given that pregnant rhesus macaques studies are labor intensive and costly, we therefore explored whether fitness advantages conferred by I1404 may be specific to pregnancy using pregnant mice.

**Figure 2.**
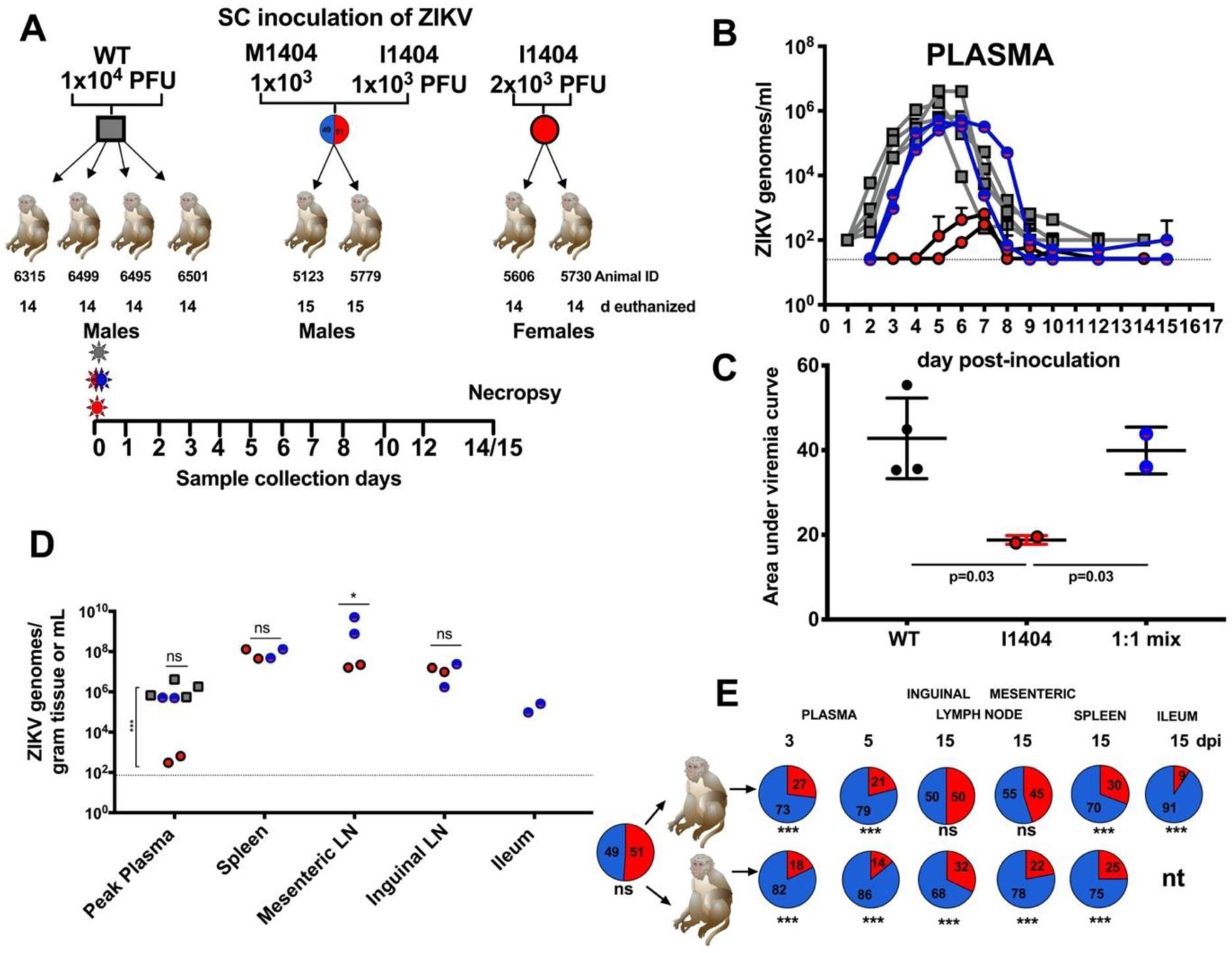
A ZIKV mutant, I1404, generates lower viremias and is less abundant than M1404 in tissues of non-pregnant rhesus macaques after mixed inoculation. **(A**) Experimental design for subcutaneous (SC) ZIKV inoculation of male and non-pregnant female rhesus macaques with ZIKV WT (left), a 1:1 mixture of infectious clone derived ZIKV I1404 and M1404 (middle) or infectious clone derived ZIKV I1404 (right) at indicated doses. The 1:1 mixture was verified by sequencing the inoculum prior to administration to macaques. Blood was collected daily from 1 to 8, 10 and 12, and at either 14 or 15 dpi, at which point animals were euthanized for tissue collection. **(B)** Plasma viremia kinetics for individual rhesus macaques inoculated with ZIKV WT (grey), the 1:1 mixture (blue) or ZIKV I1404 (red). The dotted line shows the limit of detection, 1.4×10^1^ genome copies/mL plasma. Error bars show standard deviations for triplicate measurements. **(C)** The area under the curve (AUC) for rhesus macaques infected with I1404 compared to WT or the 1:1 mix. AUC statistics were performed on log-transformed viremia measurements (one-way ANOVA)**. (D)** ZIKV RNA levels in tissues of rhesus macaques. LN is lymph node. * p=0.04; ns is not significantly different at p=0.05 (one-way ANOVA). The dotted line represents the mean limit of detection, 1.9×10^1^ ZIKV genome copies/gram tissue or mL**. (E)** Targeted quantitative sequencing flanking ZIKV 1404 showing the relative abundance of M1404 (blue) or I1404 (red) in indicated tissues from male rhesus macaques inoculated with the 1:1 mixture (*** p<0.01, ns is not significantly different at p=0.05, chi-squared tests). **Supplemental Table 3** shows the depth of sequencing coverage for the data represented here. Each dot in panels B,C and D shows the mean of triplicate qRT-PCR ZIKV RNA measurements.

### ZIKV I1404 confers fetal infection in pregnant CD-1 mice

To assess whether I1404 provides a fitness benefit in pregnancy, we inoculated pregnant CD-1 mice intraperitoneally, similar to an established model (53), with ZIKV WT, M1404, I1404, or diluent, and compared ZIKV RNA levels in maternal and fetal tissues and fetal weights and resorption rates (**Supplemental Figure 2**). Dams were euthanized on gestation day 13 (E13, where full term in mice is E21) in experiment 1 and on E13 or E19 in experiment 2 (**Figure 3A**). Back titration of residual inocula (**Figure 3B**) showed that mice were administered similar RNA levels of ZIKV M1404 and I1404 for both experiments. The WT inoculum (experiment 1 only) was lower. No statistically significant differences in rates of fetal resorption or fetal weight were observed in either experiment or at either gestation day (p>0.05, Fishers’ exact tests for resorption rates, p>0.05 for mean weights compared with ANOVA multiple comparisons) (**Supplemental Figure 2A-D**). On E13, mean maternal spleen ZIKV RNA levels for WT were significantly higher than for M1404-inoculated mice (p=0.02, two-way ANOVA) (**Figure 3C**) but no significant differences in rates of ZIKV detection or mean RNA levels in placentas were observed across the 3 ZIKV inoculated groups (p>0.05, two-way ANOVA) (**Figure 3D**). Despite similar ZIKV RNA levels in placentas, ZIKV RNA was only detected in fetuses in the ZIKV 1404I group (1404I : 9/43 [20%] versus 1404M : 0/43 [0%], p=0.003, Fisher’s exact test) (**Figure 3E**). These fetuses were from 7 different mothers. Sanger sequencing from 3 ZIKV RNA positive fetuses from different mothers in the I1404-inoculated group at E13 showed retention of I1404 in 1 fetus and reversion to M1404 in the other 2; for one mother/fetus pair the reversion was detected only in the fetus and not in the maternal spleen or placenta (**Figure 3F**). At E19, the magnitude of mean ZIKV RNA levels in spleens and the rates and magnitude of mean ZIKV RNA levels in placentas were significantly higher in mice infected with I1404 than M1404 (levels: p<0.01 for spleen, p<0.001 for placenta, one-way ANOVA, placenta infection rates: I1404: 10/20 [50%] versus M1404: 2/20 [10%], p=0.01, Fisher’s exact test) (**Figure 3G-H**). Only 1 E19 fetus head but no fetal bodies in the I1404 group tested ZIKV RNA positive (**Figure 3 I-J**) (rates not significantly different, Fisher’s exact test). Depending on the time of tissue collection, these data show that I1404 confers a fitness advantage in pregnancy by increasing the magnitude and rate of feto-placental infection during murine gestation.

**Figure 3.**
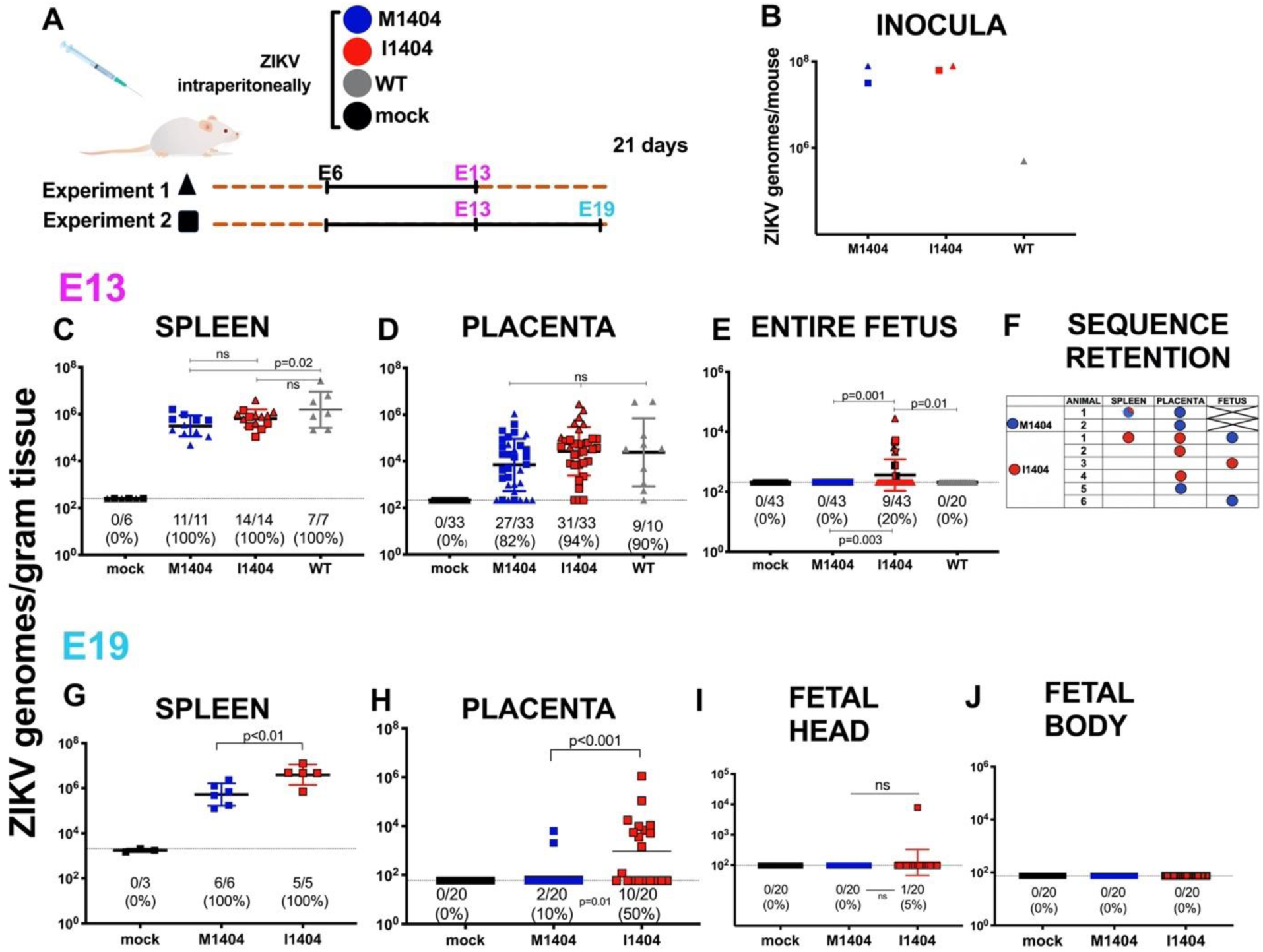
ZIKV I1404 produces higher spleen and placental ZIKV RNA titers and confers fetal infection in infected pregnant mice. **(A)** Experimental design showing intraperitoneal inoculation of pregnant CD-1 mice with ZIKV M1404 (blue), I1404 (red), WT (grey), or mock (DMEM, black) on embryonic day 6 (E6) of pregnancy, where full term in mice is E21. Two experiments were performed. In experiment 1 (triangles), dams were euthanized on E13. In experiment 2 (squares), dams were euthanized on E13 or E19. (**B)** Back titration of residual inocula. ZIKV RNA levels in maternal spleens **(C)**, placentas **(D)** and **(E)** fetuses at E13. Different patterns of shading for triangles and squares in E show different dams (N=7) from which infected fetuses were detected. (**F**) Sanger sequencing of maternal tissues or fetuses shows I1404 reverts to M1404 in some animals. An ‘X’ indicates no fetuses had detectable ZIKV RNA so could not be sequenced. An empty field indicates sequencing was not attempted from that sample. ZIKV RNA levels in (**G)** maternal spleens **(H)** placentas, **(I)** fetus heads, and **(J)** fetus bodies at E19. Each dot represents one tissue sample and is reported as the mean of 3 ZIKV RNA qRT-PCR replicates. Group means are shown as black lines, and include samples with no detectable ZIKV RNA, which were reported at the limit of detection (LOD). Only samples with 3/3 qRT-PCR replicates with detectable ZIKV RNA are reported above the LOD. The dotted line denotes the LOD, which was a mean of 1.8×10^1^ ZIKV RNA copies/gram tissue for all panels except G, where the LOD was 3.3×10^3^ ZIKV RNA copies/gram tissue. Statistical analyses comparing means used ANOVA multiple comparisons. Rates of ZIKV RNA positive samples in each group were compared with Fisher’s exact statistics.

### ZIKV I1404 mutant is more poorly transmitted than M1404 by *Aedes aegypti*

Given that fitness modifying mutations evolved by arboviruses in one host must necessarily be maintained in the alternate host for the virus to persist via arthropod-borne cycling in nature, we next considered whether ZIKV I1404 affects transmission by the primary vector, *Ae. aegypti*. Mosquitoes were presented to viremic *Ifnar-/-* mice infected with ZIKV M1404 or I1404 (**Figure 4A**). Engorged mosquitoes were held for 7 days after ingesting blood from mice that had matched ZIKV RNA levels (**Figure 4B**) and then dissected and assayed to measure rates and levels of ZIKV RNA in bodies (for infection), legs and wings (disseminated infection), and saliva (a proxy for transmission). The 7-day incubation period was chosen since it represents the time at which ZIKV exposed mosquitoes exhibit maximal infection rates (54). All *Ae. aegypti* that ingested viremic mouse blood became infected and mean ZIKV RNA levels between groups of mosquito bodies were not different (not significant, p>0.05, unpaired t-test) (**Figure 4C**). Although all mosquitoes also developed disseminated infections in legs and wings, the mean ZIKV RNA level was significantly higher for the I1404 cohort compared to M1404 cohort (p<0.007, unpaired t-test) (**Figure 4D**). Despite higher dissemination titers, *Ae. aegypti* transmitted ZIKV I1404 more poorly than M1404 (1404I: 3/20 [15%] versus M1404: 13/20 [65%], p=0.003, Fisher’s exact test) (**Figure 4E**). Sanger sequencing of three ZIKV saliva samples from mosquitoes in each group showed retention of the amino acid at 1404 (data not shown). This mosquito data revealed that ZIKV I1404 is less transmissible than M1404 in the primary vector.

**Figure 4.**
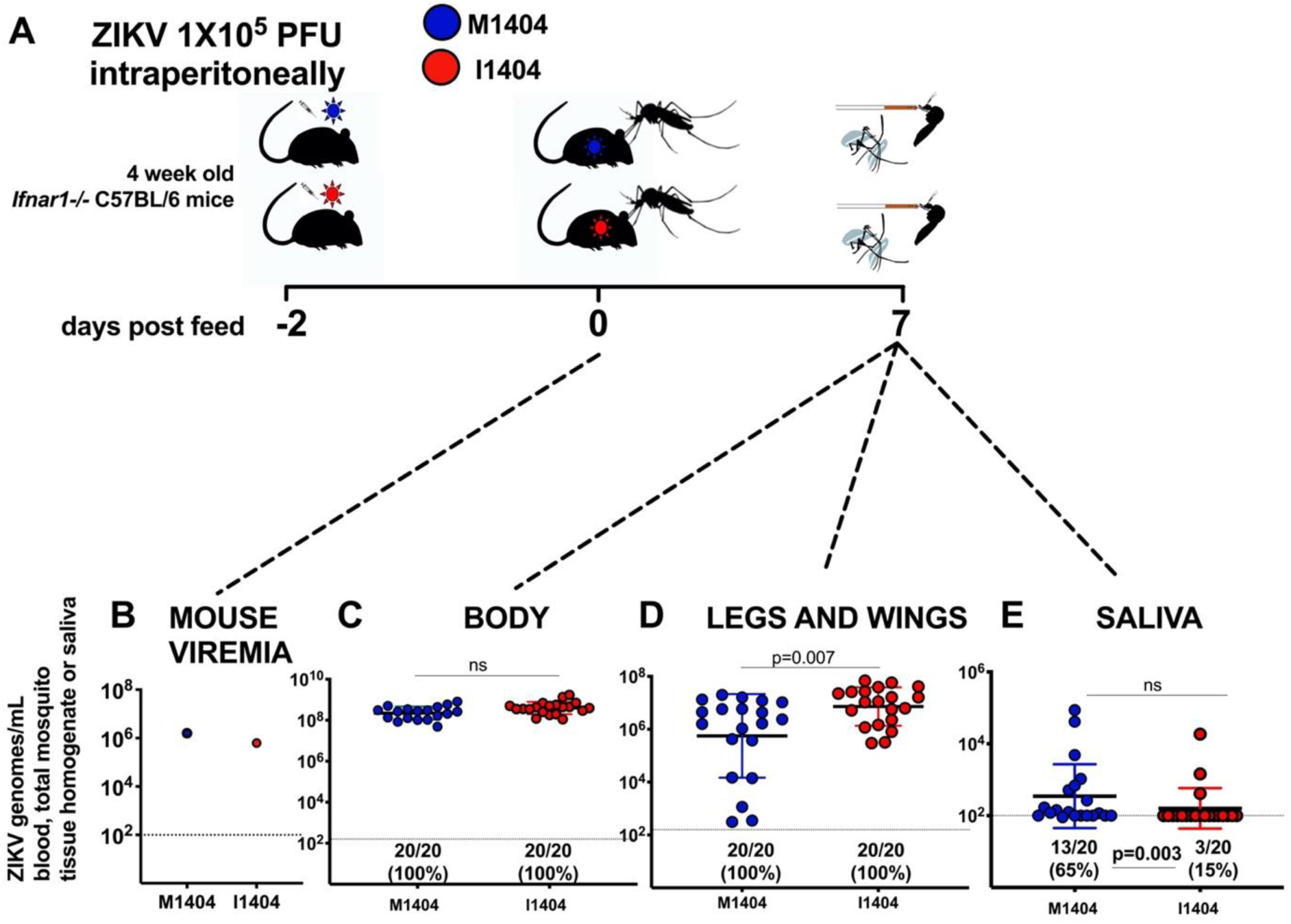
*Aedes aegypti* transmit ZIKV I1404 less efficiently than M1404. **(A)** Experimental design showing intraperitoneal inoculation of 1×10^5^ PFU per *Ifnar-/-* mouse with ZIKV M1404 (blue) or I1404 (red), two days prior to mosquito feeding, followed by presentation of viremic mice to mosquitoes, a seven day incubation period and harvesting of mosquito bodies, legs and wings, and saliva to assess infection, dissemination, and transmission rates and magnitudes of ZIKV RNA, which were quantified by qRT-PCR. **(B)** Mouse viremias immediately post-feed show that mosquitoes in both groups were exposed to similar quantities of virus in ingested blood. **(C)** ZIKV I1404 infects *Ae. aegypti* bodies and **(D)** disseminates into legs and wings at similar rates and significantly higher mean ZIKV RNA levels than M1404 but **(E)** is significantly less transmissible in saliva although transmitted doses are not different. Each dot represents mean ZIKV genomes measured in mouse blood or individual *Ae. aegypti* tissue or saliva sample 7 days post feed. Only samples with 3/3 replicates with a detectable qRT-PCR value are reported above the limit of detection. The dotted line represents the limit of detection, 2×10^2^ ZIKV genome copies/mosquito sample or blood. P values comparing mean genome levels are from unpaired t-tests. Rates were compared with Fisher’s exact statistics.

## DISCUSSION

Understanding factors that influence disease and transmissibility can lead to approaches to control ZIKV. We identified a ZIKV mutation, M1404I, that arose *de novo* or increased in frequency in experimentally inoculated pregnant rhesus macaques. Repeated detection of the mutation in tissues of multiple pregnant rhesus macaques at higher frequency than in the inoculum suggests it confers a selective advantage intrahost in pregnancy. Our experiments in mice confirm that I1404 increases ZIKV fitness in pregnancy by augmenting the magnitude and rate of placental infection and by conferring fetal infection in pregnant CD-1 mice inoculated intraperitoneally in the first trimester. By contrast, our studies in non-pregnant rhesus macaques show that I1404 is less fit than M1404, producing significantly lower viremias and decreased frequency in tissues starting from an equal ratio via mixed inoculation. The observation of inferior fitness of I1404 compared to M1404 in non-pregnant rhesus macaques parallels its low frequency in non-pregnant humans where only 5 of 543 (<1%) of publicly available ZIKV consensus genomes as of August 2020 (https://nextstrain.org/zika?c=gt-NS2B_32) possess the I1404 allele (1 of the 5 was from a microcephalic fetus). We also observed that ZIKV I1404 is not as efficiently transmitted by *Ae. aegypti*, which may further explain its low frequency in recent ZIKV outbreaks where the most common transmission route was human-mosquito-human.

The kinetics of and mechanisms by which I1404 increases ZIKV infection of placental and fetal tissues in murine pregnancy merit further study. Although I1404 enhanced feto-placental infection, reversion to wild type M1404 in some infected fetuses on at gestation day E13 suggests that M is the preferred amino acid in some tissues. Detection of ZIKV RNA in 20% of I1404 fetuses at E13 but only 5% of fetal heads and no fetal bodies on E19 also raises new questions. The disparity in fetal infection rate between E13 and E19 is likely not related to the pregnant mice deriving from different cohorts since the same rates were observed across the E13 fetuses from both experiments where the cohorts of mothers were different. Clearance or reduction in fetal infection below the limit of detection of our qRT-PCR assays between E13 and E19 is a possibility, and may reflect a difference in timing of maximum ZIKV levels, where fetal infection may peak earlier than placental infection. Defining the kinetics of fetal or placental ZIKV RNA levels over time is difficult to assess experimentally since evaluating infection involves destroying the fetus or placenta. It is also possible that since ZIKV I1404 infects fetuses at low rates, with the variable rate from experiment 1 to 2 representing a stochastic effect. ZIKV infects many placental cell types including trophoblasts, endothelial cells, fibroblasts, and fetal macrophages (55, 56), as well as multiple additional fetal cell types (24, 56–59). ZIKV I1404 may also confer infection of certain of these cell targets or augment escape from their antiviral responses in ways that M1404 cannot.

The 1404 locus is in the NS2B coding region, which encodes a 130 amino acid protein that acts as a co-factor for NS3, the protease. Relative to other flaviviral proteins, the function(s) of NS2B are poorly understood. NS2B consists of 3 transmembrane domains (TMD) (60, 61). Mutations in the NS2B TMD decrease yellow fever virus replication (62) and can modify virus assembly of Japanese encephalitis virus (60). The 1404 mutant identified in this study is located within NS2B TMD pass 2. Nuclear magnetic resonance of dengue virus NS2B indicates that TMD-TMD interactions might promote membrane fluidity or facilitate interactions with other flavivirus proteins (61). Future studies could focus on sub-cellular changes in virus-virus or virus-cell interactions mediated by M1404I that might impact cell tropism, infectivity, and immune responses.

Here we employed experimental infection of non-pregnant macaques, pregnant mice, and vector mosquitoes to study fitness of a ZIKV mutation we initially identified in pregnant rhesus macaques (**Figure 5**). The data from this study support the idea that viruses do not necessarily evolve to become more infectious or virulent, especially if those traits reduce transmissibility. Despite increasing in frequency in pregnant macaques and conferring infection of fetal mice, our data show that the I1404 mutant identified in pregnant rhesus macaques is less transmissible by vectors, as measured by lower ZIKV RNA transmission rates in saliva capture assays, and also less capable of generating viremias in non-pregnant rhesus macaques sufficient to infect feeding vectors. Although higher levels or longer periods of viremia in pregnant macaques may result from feto-placental ‘spill-back’ to the mother, a pattern we anecdotally observed in viremic pregnant rhesus macaques whose fetuses died and were removed via fetectomy and then became aviremic several days later, fetal infection is generally considered a transmission ‘dead-end’ for arboviruses. As such, arboviral mutations that augment fetal infection also need to be neutral or fitness-enhancing in mosquitoes to persist in human-mosquito-human cycling. Since we observed decreased fitness in vectors, the M1404I mutation identified in this study is not likely to spread in human-mosquito-human transmission in ZIKV outbreaks. A unique feature of this study was use of two vertebrate models of ZIKV disease as well as vector competence assays. The combined data from these three systems underscores the importance of investigating consequences of arboviral mutation in both vertebrates and invertebrates, to fully understand their roles in outbreak spread (63).

**Figure 5:**
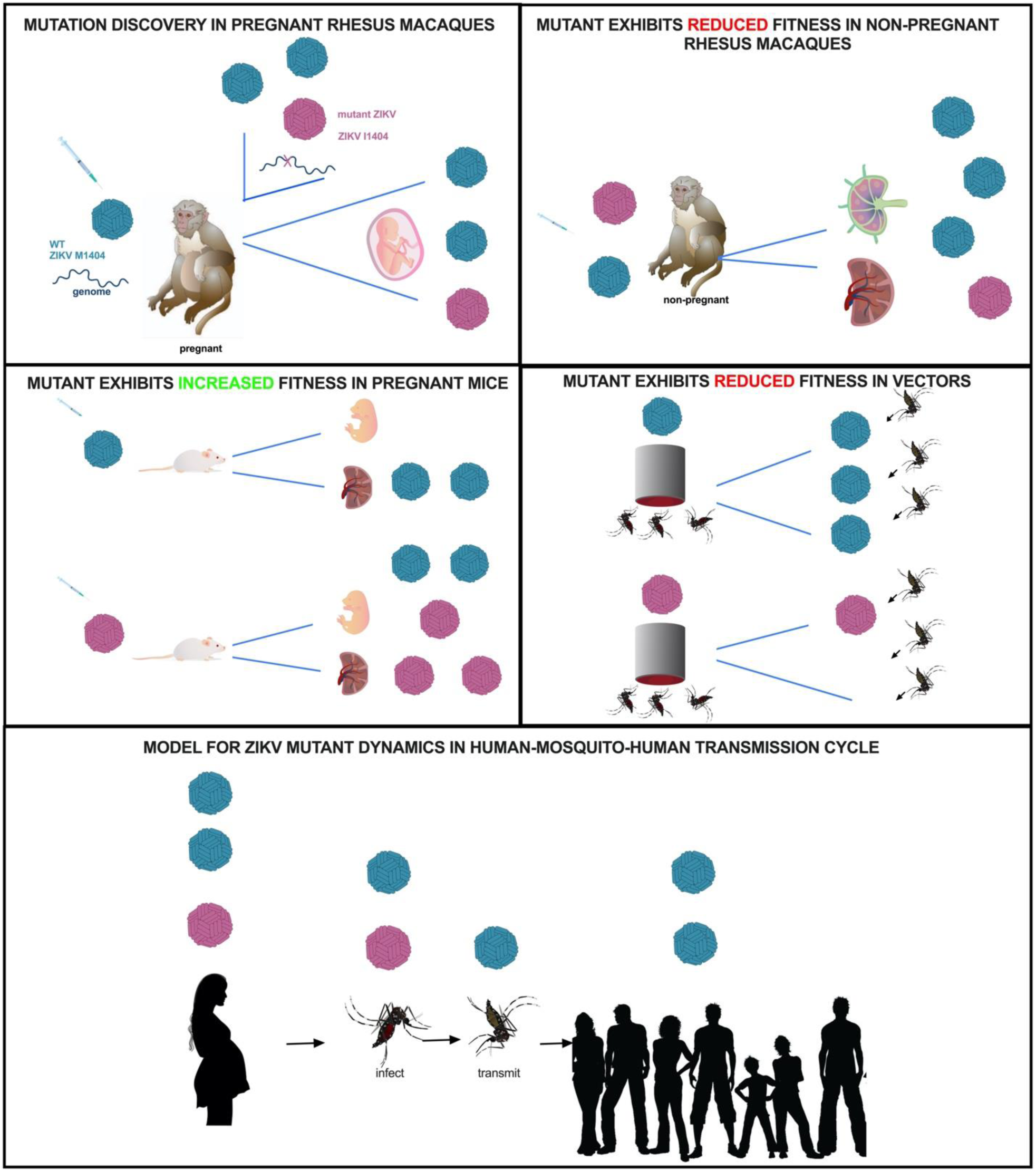
Fitness dynamics for mutant ZIKV. Visual representation of ZIKV M1404 (blue) and I1404 (red) fitness dynamics in the experimental systems used in this study. The bottom panel shows a model for possible transmission dynamics of the mutant in human-mosquito-human cycling, where human infection dynamics are predicted from observations in rhesus macaques.

## MATERIALS AND METHODS

### Rhesus macaques

Details for the studies with pregnant rhesus macaques are described elsewhere (24, 34). For non-pregnant animals, healthy male or female rhesus macaques (*Macaca mulatta*) were used in this study. All rhesus macaques were born at the California National Primate Research Center (CNPRC). Animals #5123, 5779, 5606, and 5730 received an HIV envelope protein as part of another study, but were never challenged with HIV. Prior to ZIKV inoculation, animals were housed indoors in stainless steel cages, and exposed to a 12h light/dark cycle, 18-23°C, and 30-70% room humidity. Rhesus macaques were provided with water *ad libitum* and received commercial chow a high protein diet commercial chow and fresh fruit supplements. Macaques were observed at least twice daily for clinical signs of disease including inappetence, stool quality, dehydration, diarrhea, and lethargy, and were given supportive care (including nutritional supplements) as needed. Clinical signs were rare and mild.

### Mice

Timed pregnant CD-1 mice were purchased from Charles River Laboratories (Sacramento, CA). Animals were housed in a BSL-3 facility at University of California, Davis, prior to any procedure. A maximum of 4 dams were caged together at each time, with 12 hour light/dark cycle, 18-23°C, 30-70% room humidity, social enhancers and access to mouse chow and water *ad libitum*. Non-pregnant 2 month old female Ifnar1 (IFN-α/βR−/−; C57BL/6, B6.129S2-Ifnar1tm1Agt/Mmjax, The Jackson Laboratory) were used as bloodmeal sources for vector competence studies. Mice were anesthetized prior to mosquito exposure with a mixture of ketamine (VETone Zetamine CIII, 75 mg/kg), xylazine (AnaSed, 10 mg/kg), and acepromazine (AceproJect, 1 mg/kg) solution administered intraperitoneally. Immediately after mosquito feeds, mice were euthanized while still under anesthesia via exsanguination using cardiac puncture followed by cervical dislocation.

### Animal Use

The University of California, Davis is accredited by the Association for Assessment and Accreditation Laboratory Animal Care International (AAALAC). Animal care was performed in compliance with the 2011 Guide for the Care and Use of Laboratory Animals provided by the Institute for Laboratory Animal Research and both rhesus macaque and mouse studies were approved by the Institutional Animal Care and Use Committee (IACUC) of the University of California, Davis. All mouse procedures were approved under protocol #19404. All rhesus macaque procedures were approved under protocols #19211 and 19695.

### Mosquitoes

The fourth generation of *Aedes aegypti* originally collected in 2015 in Puerto Rico were used in this study. Mosquitoes were maintained in a colony in an insectary at the University of California, Davis. Twenty four hours prior to exposure to a mouse, mosquitoes were transferred into pint cartons and transported into to a BSL-3 facility to acclimate in a humidified chamber set to a 12 hour light/dark cycle, 27°C, 80% humidity. After ingestion of blood from viremia ZIKV infected mice, mosquitoes were presented with 10% sucrose *ad libitum.*

### Cell lines

African green monkey kidney cells (Vero; ATCC CCL-81) were cultured at 37°C in 5% CO_2_ cultured in Dulbecco’s Modified Eagle Medium (DMEM, Gibco, Thermo Fisher Scientific) supplemented with 2% fetal bovine serum FBS (Gibco, Thermo Fisher Scientific) and 1% penicillin/streptomycin (Gibco, Thermo Fisher Scientific). Baby hamster kidney cells (BHK21; ATCC CCL-10) were cultured in the same conditions as Vero cells but supplemented with 10% FBS. *Aedes albopictus* cells (C6/36; ATCC CRL-1660) were cultured in Schneider’s insect medium (Caisson Labs) supplemented with 20% FBS and 1% P/S at 28°C and atmospheric CO_2_.

### Viruses

#### Wild type Zika virus stock

For growth curves and *in vivo* experiments in pregnant and non-pregnant rhesus macaque and pregnant mice, the 2015 Brazilian ZIKV strain SPH2015 (Genbank accession number KU321639) was used. This virus was originally isolated from a human transfusion recipient in São Paulo, Brazil and then passaged 3 times in Vero cells. We refer to this strain as wildtype (WT) throughout the paper. A 2015 Puerto Rico ZIKV strain PRVABC59 (Genbank accession number KU501215.1, Vero passage 4) was also used in pregnant rhesus macaque that were sequenced for this project. We refer to this strain as ‘ZIKV Puerto Rico 2015’ throughout the paper.

#### Generation of Infectious clone derived M1404 and I1404 Zika viruses

To focus on the 1404 locus as a determinant of phenotype, we generated 2 infectious clones that were identical in sequence except for at 1404. To start, we modified a ZIKV infectious clone made from the sequence of a 2015 Brazilian ZIKV strain, Paraiba_01/2015 (16) to match the amino acid sequence of WT (SPH2015), which encodes M1404. Six mutations were inserted into the Pariaba_01/2015 clone at positions V313I, Y916H, V1143M, H1857Y, I2295M and I2445M of the ZIKV polyprotein to generate the WT amino acid sequence of strain SPH2015. We refer to this clone as ‘ZIKV M1404’. Next the M1404 clone was mutated to change the amino acid at 1404 from G>A (AUG [methionine] > AUA [isoleucine]) to generate the I1404 clone. The ZIKV I1404 clone is identical to the M1404 clone except at NS2B locus 1404. Each of the mutagenized loci were verified by Sanger sequencing. Infectious viruses were rescued from ZIKV M1404 and I1404 clones by electroporating plasmid DNA into BHK cells. 2.5μg of plasmid DNA was electroporated at 110V, 1750uF capacitance, and no resistance using an ECM 630 electro cell manipulator (BTX Harvard Apparatus) into ≈50% confluent T75 flasks of BHK21 cells. Cells were then centrifuged for 5 minutes at 1500 revolutions per minute (RPM) and resuspended in DMEM in T25 flasks for 3 days of incubation at 37°C with 5% CO_2_. After 3 days, the supernatant was collected, spun at 1500 RPM for 5 minutes to remove cell debris, and stored at -80°C in 400μL aliquots. These recovered infectious clone derived viruses, termed ZIKV M1404 and ZIKV I1404 were used in growth curves, mosquito, pregnant mouse and non-pregnant rhesus macaque experiments. The genotypic integrity of both infectious clone derived viruses was verified by whole-genome Sanger sequencing from the electroporation-harvested stocks.

#### Zika virus titrations using plaque assays

Infectious ZIKV from electroporation-rescued stocks of infectious clone-derived viruses, inocula, growth curves, mosquito saliva, and RT-qPCR positive mouse fetuses were titrated using plaque assays. Plaque assays were performed in confluent 6-well plates of Vero cells that were inoculated with 250μL of ten-fold dilutions of virus stock or sample resuspended in 2% FBS DMEM. Cells were incubated for 1 hour at 37°C and 5% CO_2_ and plates were rocked every 15 minutes to prevent cell death due to desiccation. After 1 hour, 3 mL of 0.5% agarose mixed with 2% FBS/DMEM was added to each well to generate a solid agar plug. The cells were then incubated for 7 days at 37°C and 5% CO_2._ After 7 days, the cells were fixed with 4% formalin for 30 minutes, the agar plugs were removed, and the cells were stained with 0.025% crystal violet in 20% ethanol in order to visualize and quantify plaques. Samples were tested in duplicate and the average titer is reported as the number of plaques visible against a white background. The limit of detection was 40 plaque-forming units (PFU).

#### Zika virus RNA isolation

ZIKV RNA was isolated using MagMax (Thermo Fisher) or Qiazol (Qiagen). The MagMax system was used to extract ZIKV RNA from growth curve supernatants, rhesus macaque plasma, and homogenized mosquito bodies, legs and wings, and saliva. The MagMax Viral RNA Extraction Kit was used according to manufacturer’s recommendations. For cell supernatant and mosquito homogenates, a total of 100 μL of sample was extracted and for rhesus macaque plasma the volume extracted was 300 μL. Maternal spleens, placentas and fetuses from the mouse experiments and solid tissues from rhesus macaque were extracted using Qiazol. Tissues stored in RNAlater were first removed from that solution prior to RNA extraction. Using clean forceps and scissors, tissues were removed from RNAlater, and a portion between 20-50 mg was cut and placed in a pre-weighed tube containing a 0.5 mm glass ball bearing (Fisher). Tubes were then re-weighed and 900 uL of Qiazol solution was added before trituration in a TissueLyzer (Retsch) machine. Tissues were homogenized for 2 m at 30 shakes/second (s). If liquefaction of tissues was incomplete, samples were homogenized for an additional 2 m at 30 shakes/s. The homogenate was centrifuged for 2 m at 14,000 g to clarify the supernatant, which was tested. Viral RNA from homogenates were extracted following the manufacturer’s kit instructions. The Qiazol protocol was modified for the E19 fetuses since they were large. E19 fetus samples were homogenized in 1 mL of DMEM and 200 μL of the homogenate was added to 900 μL of lysis reagent, after which the protocol followed the manufacturer’s instructions. All RNA extracts were eluted in 60μL of Qiagen elution buffer and archived at -80°C until further analysis.

#### Zika virus RNA quantification by qRT-PCR

ZIKV RNA samples were each measured in triplicate on an Applied Biosystems ViiA 7 machine using a Taqman Fast Virus 1-Step MasterMix (Thermo Fisher) with primers ZIKV 1087 forward (CCGCTGCCCAACACAAG), ZIKV 1163c reverse (CCACTAACGTTCTTTTGCAGACAT), and ZIKV 1108-FAM probe (AGCCTACCTTGACAAGCAGTCAGACACTCAA) according to the protocol in Lanciotti *et al.* (64). The protocol was modified by increasing the initial volume of sample tested to 9.6 μL to increase sensitivity. Samples were only considered positive if all three replicates yielded at detectable cycle threshold (Ct) value of less than the cutoff of the assay, 40. For each 96-well plate where samples were tested, a standard curve was generated from serial dilutions of a synthetic DNA of known concentration corresponding to the qRT-PCR target region. The reported limit of detection (LOD) on each graph shows the mean of all samples with a detectable Ct that are included in the graph. Where means are reported for a group of measurements, samples with no detectable ZIKV RNA were included in measurements where their values were reported at the LOD.

#### In vitro Zika virus growth assays

Vero and C6/36 cells were inoculated in triplicate with ZIKV M1404, I1404 or WT at a multiplicity of infection (MOI) of 0.01 (Vero only) or 0.5. Cells in one well in the plate were counted immediately prior to infection and the MOI was adjusted according to the cell count. The cells were inoculated by overlaying 150 μL of virus for 1 hour at 37°C in 5% CO_2_ for Vero and 28°C in ambient CO_2_ for C6/36 with gentle rocking every 15 minutes. After 1 hour, the cells were washed three times with phosphate buffered saline (PBS) to remove residual unbound viruses, and 2 mL/well of DMEM was added. At time points 0, 12, 24, 48, 72, and, for Vero, 96 hpi (hours post inoculation) 160 μL/well was collected and archived at -80°C until it was tested in triplicate to determine ZIKV RNA levels by qRT-PCR. Data shown are the percent of the mean 0 hpi residual input ZIKV RNA in genomes/ml. Each data point shows the mean measurement from 3 wells that were each measured in triplicate by qRT-PCR.

#### Experimental inoculation of non-pregnant rhesus macaques with Zika virus

Studies with pregnant rhesus macaques were described elsewhere [23,33]. Non-pregnant female and male rhesus macaques were inoculated subcutaneously or intravenously with 1 mL of ZIKV WT at 1×10^4^ PFU, a 1:1 mixture of ZIKV M1404 and I1404, where 1×10^3^ PFU of each virus was in the inoculum, or 2×10^3^ PFU of ZIKV I1404. All inocula were back-titrated immediately after inoculation without freezing using plaque assays to verify the administered doses. All inocula were re-sequenced to verify identity at the 1404 locus. The mixed inoculum was also checked prior to inoculation by next generation sequencing flanking the 1404 locus and was verified to contain 49% M1404 and 51% I1404. Urine and blood were collected from rhesus macaques daily for 7 days and then every other day until 14 dpi. Macaques were anesthetized with ketamine hydrochloride (10 mg/kg) and samples were processed according to previously described methodologies (34). At 14 dpi, rhesus macaques were euthanized and necropsied. During necropsies, tissues were grossly evaluated *in situ*, and then excised using forceps and then dissected with disposable razor blades to minimize cross-contamination. Tissues were either snap frozen by immersion in liquid nitrogen or stored in RNAlater solution. RNAlater samples were held at 4°C for 24 hours then transferred to -80°C for further analysis.

#### Experimental inoculation of pregnant mice with Zika virus

Two ZIKV experiments were performed with pregnant mice. Numbers of mice are shown in figures. For both experiments, mice were sedated with isoflurane and inoculated intraperitoneally with 100 μL of 1×10^5^ PFU of ZIKV WT, M1404, I1404, or DMEM on gestation day 6 (E6). Inocula doses were verified by qRT-PCR. All mice in experiment 1 and some in experiment two were euthanized on E13, where full term is 21 days. Some mice in experiment two were euthanized later, at E19. After inoculation, mice were monitored twice daily by visual observation. On E13 or E19, mice were sedated with isoflurane and euthanized by cervical dislocation. Uterine horns were exposed and visually observed for viable or aborted/resorbed fetuses. The maternal spleen was excised and cut in half. Half was placed in a 2 mL tube containing 1 mL of RNAlater and the other half was stored in a 2 mL tube with 1 mL of DMEM. Placentas were collected in 2 mL tubes containing 1 mL of DMEM. Fetuses were weighed and their length was measured, and then they were collected in a 2 mL tube containing 1 mL of DMEM. Between each dam, forceps and scissors were immersed in 10% bleach and then 70% ethanol solution to minimize cross-contamination across animals. Samples in DMEM were immediately stored at -80°C. Samples in RNAlater were stored at 4°C for 24h and then transferred to -80°C until further analysis.

#### Experimental vector competence of Zika virus in Aedes aegypti

Ifnar1-/- C57BL/6 mice were intraperitoneally inoculated with 1×10^5^ PFU of ZIKV WT, M1404 or I1404 two days prior to presentation to female *Ae. aegypti* mosquitoes. Mice were anesthetized prior to mosquito presentation with a ketamine (75 mg/kg), xylazine (10 mg/kg) and acepromazine (1 mg/kg) solution administered intraperitoneally. Viremic mice were presented to sugar deprived mosquitoes for 1 hour. After 1 hour, mouse blood was collected to measure ZIKV RNA levels immediately after mosquito feeding. Mice were then euthanized by cervical dislocation. Engorged female mosquitoes were sorted from non-fed individuals by visual examination and then held for 7 days. On day 7, mosquitoes were cold anesthetized for 3-5 minutes at -20°C and then their legs and wings were removed with forceps while immobilized on ice. Saliva was collected by inserting the proboscis into a glass capillary tube containing FBS for 30 minutes. Mosquito bodies, legs and wings, and saliva including the capillary were placed into 2 mL tubes containing 250 μL of DMEM and a glass bead (Fisher). Samples were immediately archived at -80°C for further analysis. After thawing, mosquito samples were homogenized for 2 m at 30 shakes/second (s) in a TissueLyzer. The homogenate was centrifuged for 2 m at 14,000 g to clarify the supernatant, which was tested

#### Zika virus sequencing and sequence analyses

Plasmid sequences of infectious clones and identities of virus stocks as well as selected samples from mice and mosquitoes were verified by Sanger sequenced using primers flanking the entire genome or 1404 (**Supplemental Table 1**). Extracted RNA samples were amplified using a Qiagen One-Step RT-PCR kit and forward primer: AGCTGTTGGCCTGATATGCG with reverse primer: AGCTGCAAAGGGTATGGCTA. The cycling conditions were as follows: 50°C for 30 min, 95°C for 15 min, 40 cycles of 94°C for 1 min, 57°C for 1 min, 72°C for 1 min, followed by 72°C for 10 min and 4°C hold. Samples were sequenced at the University of California, Davis Sequencing core facility. Chromatograms were visualized and sequences were called using Sequencher (GeneCodes). The complete ZIKV genome from the inoculum and samples from pregnant rhesus macaques 5338 whose fetus died 7 dpi was sequenced at both the University of California, San Francisco using an established protocol (65) and Lawrence Livermore National Laboratories using a different established protocol (66). All other rhesus macaques and other samples in this study were sequenced using a different next generation sequencing protocol that was previously described (67, 68) and adapted here to flank 1404. After viral RNA isolation using Qiazol and RNA quantification by qRT-PCR, at least 1000 genome copies/sample were used to generate libraries for sequencing. 5 μL of ZIKV RNA was used in a cDNA synthesis reaction with a SuperScript IV kit (Invitrogen) in addition to 6 μL of nuclease free water, 1 μL of 10mM dNTP mix and 1 μL of random hexamers. The mixture was heated to 70°C for 7 minutes and placed immediately on ice. A new mixture containing 4 μL of 5x SSIV (SuperScript IV) buffer with 1 μL of 100 mM DTT, 1 μL of RNAse inhibitor and 1 μL of SSIV reverse transcriptase was added and the cDNA synthesis occurred at thermocycler conditions of 23°C for 10 min, 50°C for 45 min, 55°C for 15 min, 80°C for 10 min and 4°C until further use. Position 1404 was amplified using 1 μL each of 10 μM of forward primer: CCCTAGCGAAGTACTCACAGCT, reverse primer: TACACTCCATCTGTGGTCTCCC, 2.5 μL of cDNA, 15 μL of nuclease-free water, 0.5 μL of Q5 High Fidelity DNA Polymerase (New England Biolabs), 1 μL of 10mM dNTPs and 5 μL of 5x Q5 reaction buffer followed by 98°C for 30 s, 95°C for 15 s, 65°C for 5 min, then repetition of steps 2 and 3 for 34 additional cycles and then a hold at 4°C until further use. PCR products were purified using Agencourt Ampure XP magnetic beads (Beckman Coulter) at a 1.8:1 ratio of beads to sample. Sequencing libraries were next generated using a Kapa Hyper prep kit (Roche). Specifically, ends were repaired by mixing 1.75 μL of end-repair and A-tailing buffer, 0.75 μL of end-repair A-tailing enzyme mix, and 12.5 μL of amplified DNA followed by incubation at 20°C for 30 m, then 65°C for 30 min. For adaptor ligations, 2.5 μL of 250 nM of NEXTflex Dual-Index DNA barcodes (Bioo Scientific) were used with 15 μL of end-repair reaction product, 2.5 μL of DNA ligase and 7.5 μL of ligation buffer incubated at 20°C for 15 min. This procedure was followed by a post-ligation cleanup using Agencourt Ampure XP magnetic beads at a ratio of 0.8:1 beads to sample. The sequencing library was then amplified using 17 μL of 2X KAPA HiFi HotStart ReadyMix (Roche), 2 μL of Illumina primer mix and 15 μL of adaptor-ligated library followed by 98°C for 45 s, 98°C for 15 s, 60°C for 30 s, 72°C for 30 s, and then a repetition of steps 2-4 for 8 cycles, followed by 72°C for 1 m and 4°C until further use. Amplified samples were then cleaned using Agencourt Ampure XP magnetic beads in a ratio of 0.8:1 beads to sample. DNA library sizes were then analyzed using a BioAnalyzer DNA 1000 kit (Agilent) and the DNA concentration was quantified using Qubit High Sensitivity DNA kit (Thermo Fisher) lLibraries were diluted to 2 nM in 10 mM of TE and samples were sequenced with MiSeq (Illumina) using a paired-end approach. We used a previously described workflow [64] to determine M1404I allele frequencies. Briefly, sequence reads were trimmed to remove primer sequences and low quality base calls before they were aligned to the Zika Paraiba_01 reference genome using BWA-mem (69). Mutants over a 3% minor allele frequency were called using SAMtools mpileup (70)and were filtered according to frequency and strand biases. After sequencing, the ratios of G (encoding M1404) versus A (encoding I1404) were calculated and are represented as a percent of total sequencing depth at the locus.

#### Statistical analyses

Statistical analyses were performed using GraphPad Prism 7 (GraphPad Software). Tests used are indicated in results and figure legends. Statistical significance is denoted by P values of less than 0.05.

## DATA AVAILABILITY

Sequencing data are available in the NCBI SRA at accession number PRJNA556052.

## ACKNOWLEDGEMENTS

We acknowledge Priya Shah and Paul Luciw for productive discussions. We acknowledge pathology services at the California National Primate Research center for assisting with rhesus macaque experiments. This work was performed under the auspices of the U.S. Department of Energy by Lawrence Livermore National Laboratory under Contract DE-AC52-07NA27344.

## AUTHOR CONTRIBUTIONS

Conceptualization: D.L., L.L.C., K.K.A.V.R., N.G., K.A. Methodology: D.L., J.B.S., W.L. L.L.C., K.K.A.V.R., Investigation: D.L., J.B.S., W.L., A.S., J.W., J.U., R.I.K., G.O., S.S., N.G., M.B., J.A., J.T., K.A.T., A.G.P., L.L.C., K.K.A.V.R, C.Y.C. Writing-original draft: D.L., L.L.C. Writing-review and editing: L.L.C. D.L., J.B.S., K.K.A.V.R., K.A.T., R.I.K., N.D.G., M.B., C.Y.C. Visualization: D.L., L.L.C., K.K.A.v.R. Supervision and project administration: L.L.C.. Funding acquisition: L.L.C., K.K.A.V.R, C.C., K.A., M.B, A.G.P, C.Y.C.

## SUPPLEMENTAL FIGURES AND TABLES

**Supplemental Figure 1.**
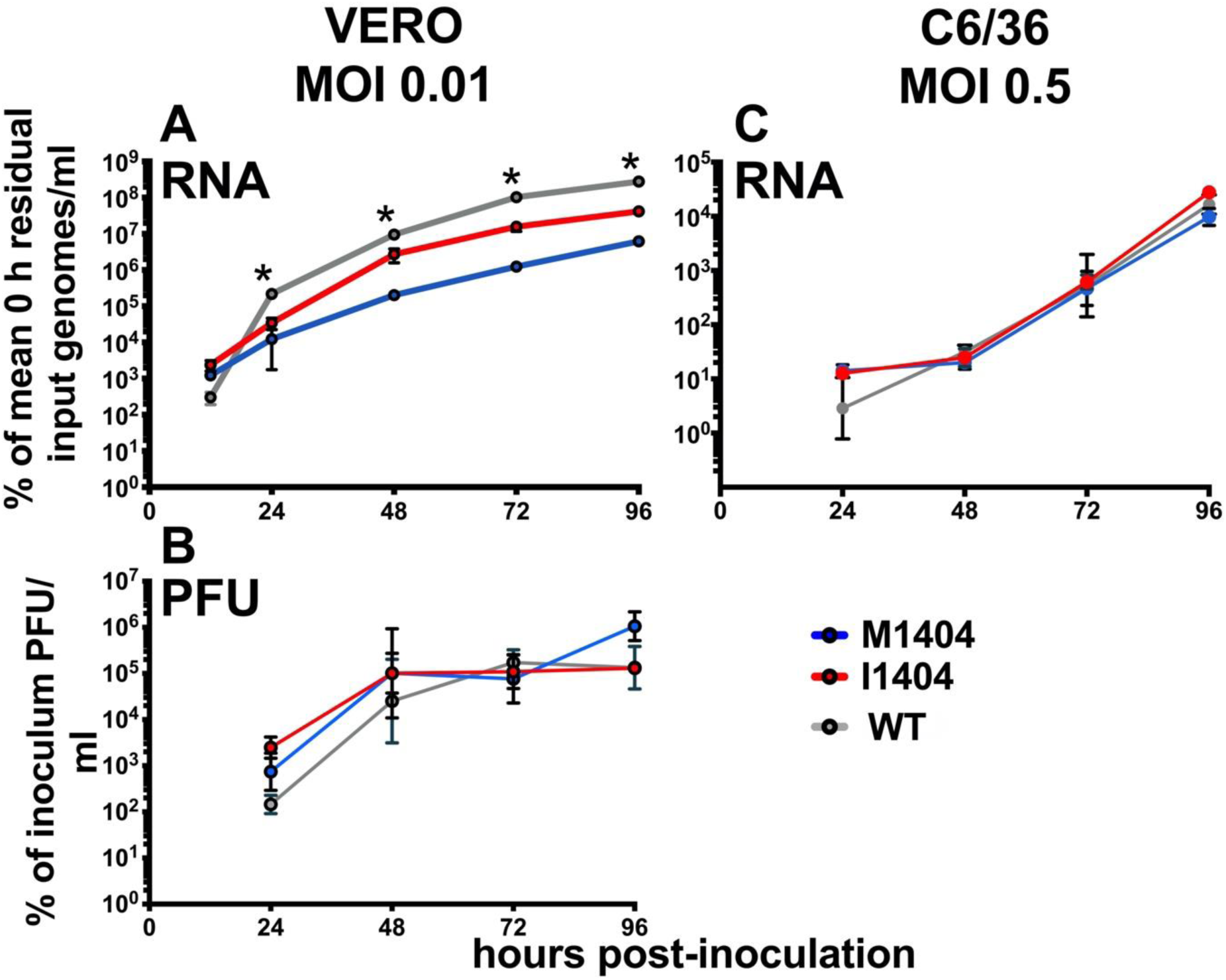
Growth kinetics of ZIKV M1404, I1404, and WT in African green monkey kidney (Vero) and *Aedes albopictus* (C6/36) cells. **(A)** ZIKV I1404 exhibits superior ZIKV RNA growth kinetics compared to M1404 from 24 to 96 hpi in Vero cells at a MOI of 0.01. Asterisks denote p<0.001 across all groups. **(B)** Infectious ZIKV levels in Vero cells from 24-96 hpi are not different between ZIKV M1404, I1404, and WT (p>0.05). **(C)** ZIKV RNA levels in C6/36 cells at a MOI of 0.5 are not different across groups (p>0.05). Each dot represents the mean of 3 replicate supernatant samples, where each supernatant was measured in 3 qRT-PCR replicates. RNA and PFU measurements for A and B were from the same supernatants. Statistical tests used were repeated measures 2-way ANOVA. Error bars show standard deviations. MOI is multiplicity of infection.

**Supplemental Figure 2.**
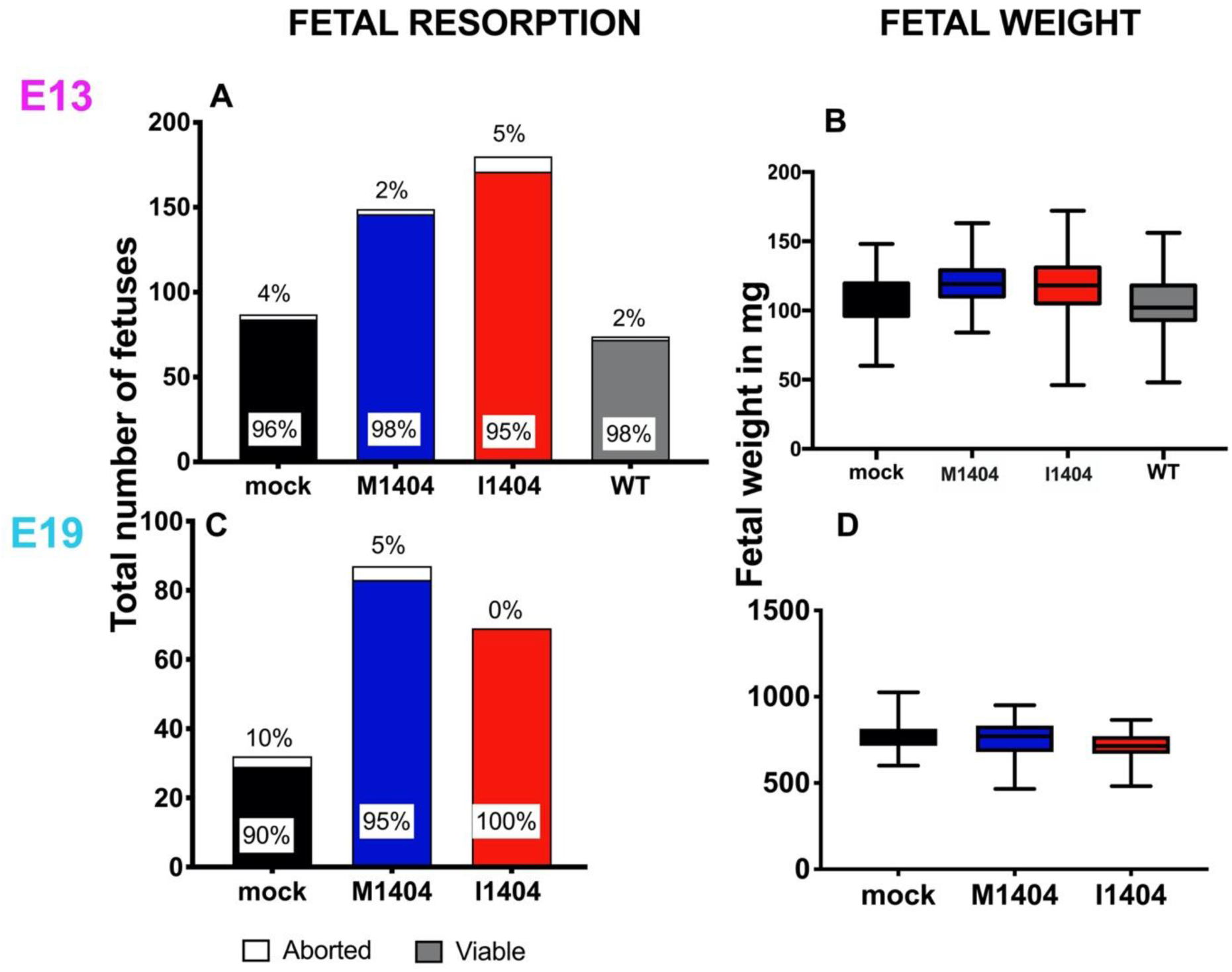
Relative to M1404 or WT, the ZIKV mutant I1404 does not augment fetal death or decrease fetal weight in ZIKV infected pregnant CD-1 mice. Mice were inoculated as shown in Figure 3A. No significant differences in rates of fetal resorption (**A**,**C**) or weight **(B**,**D)** on gestation day of harvest, E13 or E19, were detected across groups of pregnant mice infected with ZIKV M1404, I1404, WT, or that were mock-inoculated. The lines in the middle of each box for panels B and D show the mean and error bars show standard deviations. Resorption rates were compared with chi-squared statistics. Mean weights were compared with ANOVA multiple comparisons statistics.

**Supplemental Table 1.**
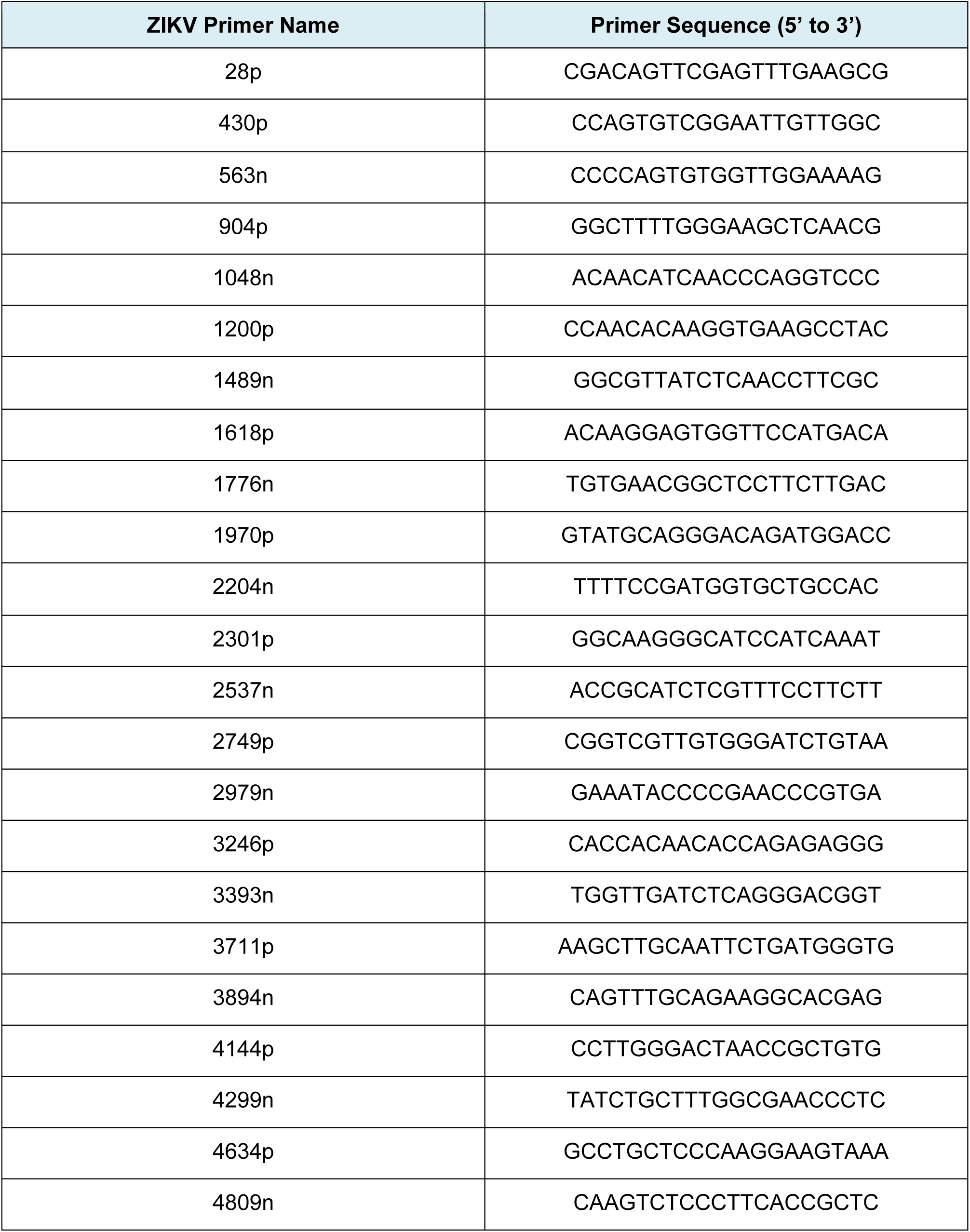
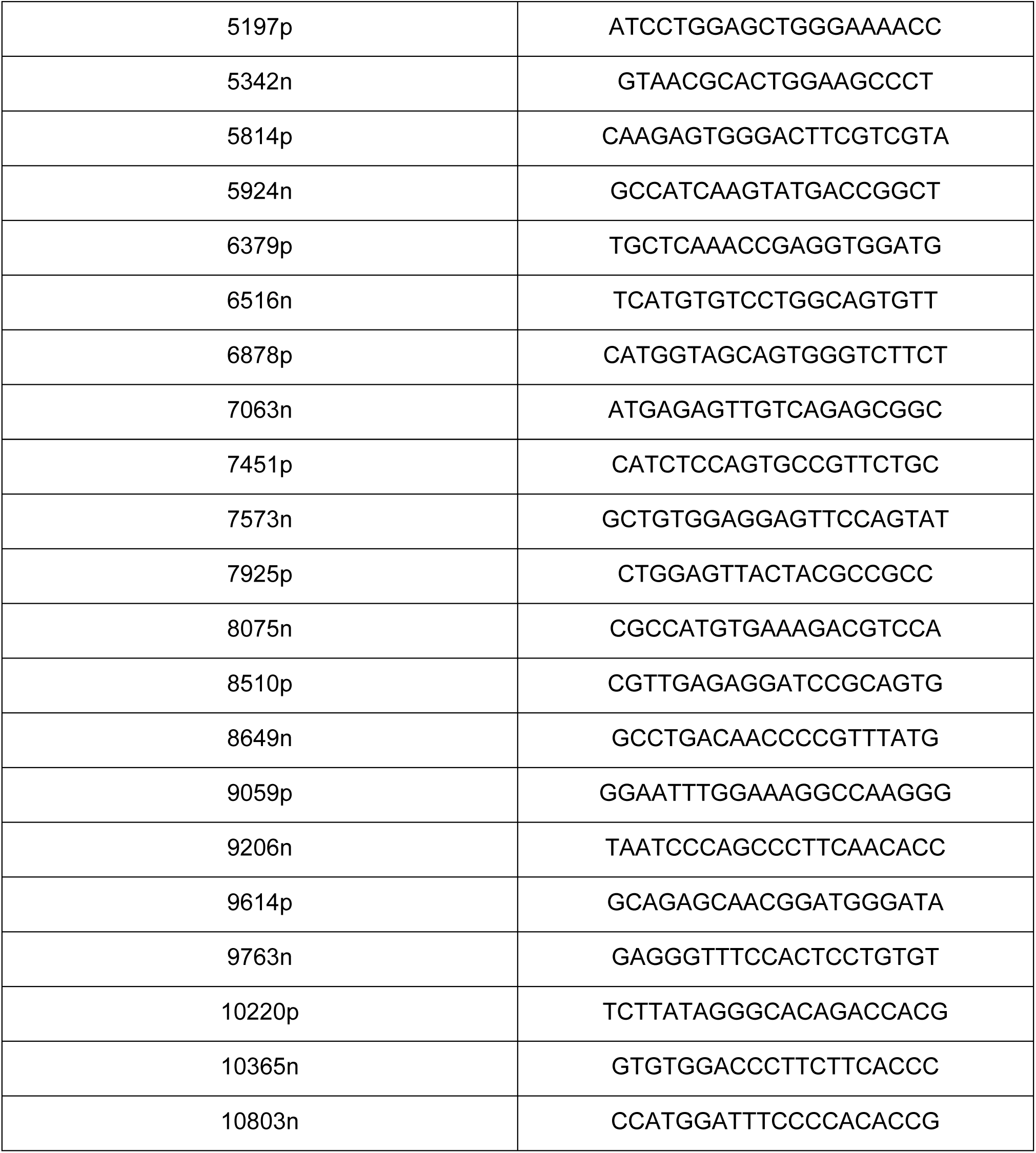
Primers used for Sanger sequencing ZIKV in this study.

**Supplemental Table 2.** Deep sequencing data showing the percentage of reads that encode ZIKV I1404 in inocula and experimentally infected pregnant rhesus macaques. For most samples, a targeted sequencing approach flanking ZIKV polyprotein amino acid 1404 was used. ^a^,^b^ and ^c^ show data from different laboratories. ^a^ is The Scripps Research Institute, ^b^ is University of California San Francisco, and ^c^ is Lawrence Livermore National Laboratories. PR is Puerto Rico, WT is SPH2015, a Brazilian strain, KU321639, N/A is not applicable, DPI is day post inoculation, IV is intravenous, IA is intraamniotic, SC is subcutaneous.

| ANIMAL IDENTIFIER or INOCULUM | ZIKV STRAIN | INOCULATION ROUTE | FETAL OUTCOME | STUDY END POINT (DPI) | TISSUE | DPI | COVERAGE AT 1404 | % I1404 MUTANT (AUA) |
| --- | --- | --- | --- | --- | --- | --- | --- | --- |
| Inoculum | PR 2015 | SC | N/A | N/A | N/A | N/A | 2242 | 0%<sup>a</sup> |
| Inoculum | WT | IV + IA | N/A | N/A | N/A | N/A | 5705 | 0.2%<sup>b</sup> |
| Inoculum | WT | IV + IA | N/A | N/A | N/A | N/A | 2296 | 1.3%<sup>a</sup> |
| 5731 | PR 2015 | SC | Near term | 125 | Maternal Spleen | 125 | 15 | 14%<sup>a</sup> |
| 5388 | WT | IV + IA | Death | 7 | Amniotic fluid | 2 | 1175 | 25%<sup>b</sup> |
| 5388 | WT | IV + IA | Death | 7 | Amniotic fluid | 7 | 2564 | 17%<sup>a</sup> |
| 5388 | WT | IV + IA | Death | 7 | Gestational Sac | 7 | 8020 | 14%<sup>a</sup> |
| 5388 | WT | IV + IA | Death | 7 | Amniotic membrane | 7 | 309 | 25%<sup>c</sup> |
| 5388 | WT | IV + IA | Death | 7 | Amniotic membrane | 7 | 4492 | 28%<sup>a</sup> |
| 5388 | WT | IV + IA | Death | 7 | Placenta full thickness | 7 | 5477 | 5%<sup>a</sup> |
| 5388 | WT | IV + IA | Death | 7 | Vagina | 7 | 777 | 19%<sup>c</sup> |
| 5388 | WT | IV + IA | Death | 7 | Spleen | 7 | 6235 | 0%<sup>a</sup> |
| 5388 | WT | IV + IA | Death | 7 | Inguinal Lymph node | 7 | 4332 | 0.1%<sup>a</sup> |
| 5388 | WT | IV + IA | Death | 7 | Fetal Lung | 7 | 6236 | 0%<sup>a</sup> |
| 5388 | WT | IV + IA | Death | 7 | Fetal Intestine | 7 | 21444 | 0%<sup>a</sup> |
| 5388 | WT | IV + IA | Death | 7 | Fetal Brain | 7 | 4581 | 0.2%<sup>a</sup> |
| 5592 | WT | IV + IA | Death | 14 | Amniotic fluid | 14 | 13 | 8%<sup>a</sup> |
| 5592 | WT | IV + IA | Death | 14 | Inguinal Lymph node | 14 | 2389 | 0%<sup>a</sup> |
| 5592 | WT | IV + IA | Death | 14 | Amniotic membrane | 14 | 7695 | 0%<sup>a</sup> |
| 5592 | WT | IV + IA | Death | 14 | Fetal intestine | 14 | 9362 | 0.1%<sup>a</sup> |
| 5592 | WT | IV + IA | Death | 14 | Fetal kidney | 14 | 6812 | 0.1%<sup>a</sup> |
| 5592 | WT | IV + IA | Death | 14 | Fetal brain | 14 | 2849 | 0.1%<sup>a</sup> |
| 4817 | WT | IV + IA | Near term | 105 | Amniotic fluid | 2 | 8000 | 0%<sup>c</sup> |
| 4817 | WT | IV + IA | Near term | 105 | Amniotic fluid | 14 | 438486 | 4%<sup>a</sup> |
| 4817 | WT | IV + IA | Near term | 105 | Amniotic fluid | 21 | 2560 | 5%<sup>a</sup> |
| 4817 | WT | IV + IA | Near term | 105 | Vagina | 105 | 1232 | 9%<sup>c</sup> |
| 5846 | WT | IV + IA | Near term | 87 | Amniotic fluid | 7 | 15869 | 13%<sup>a</sup> |
| 5846 | WT | IV + IA | Near term | 87 | Amniotic fluid | 14 | 5439 | 12%<sup>a</sup> |
| 5846 | WT | IV + IA | Near term | 87 | Amniotic fluid | 21 | 10045 | 35%<sup>a</sup> |
| 5846 | WT | IV + IA | Near term | 87 | Amniotic fluid | 28 | 333 | 2%<sup>c</sup> |
| 5846 | WT | IV + IA | Near term | 87 | Amniotic membrane | 87 | 217 | 7%<sup>c</sup> |
| 5846 | WT | IV + IA | Near term | 87 | Fetal seminal vesicle | 87 | 557 | 3%<sup>c</sup> |
| 5846 | WT | IV + IA | Near term | 87 | Urine | 40 | 1929 | 11%<sup>c</sup> |
| 5846 | WT | IV + IA | Near term | 87 | Urine | 77 | 1725 | 37%<sup>c</sup> |
| 5846 | WT | IV + IA | Near term | 87 | Placenta | 87 | 2991 | 0%<sup>c</sup> |
| 5846 | WT | IV + IA | Near term | 87 | Fetal brain | 87 | 2800 | 0%<sup>c</sup> |
| 5846 | WT | IV + IA | Near term | 87 | Amniotic fluid | 2 | 7842 | 0.3%<sup>a</sup> |
| 5207 | WT | IV + IA | Near term | 65 | Amniotic fluid | 2 | 8797 | 14%<sup>c</sup> |
| 5207 | WT | IV + IA | Near term | 65 | Placenta | 65 | 5130 | 4%<sup>c</sup> |
| 5462 | WT + PR 2015 | IV + IA | Death | 73 | Amniotic fluid | 7 | 6151 | 0%<sup>a</sup> |
| 5462 | WT | IV + IA | Death | 73 | Amniotic fluid | 14 | 6686 | 0.4%<sup>a</sup> |
| 5462 | WT | IV + IA | Death | 73 | Fetal Lung | 70 | 5016 | 0%<sup>a</sup> |
| 5507 | PR 2015 | SC | Death | 30 | Amniotic membrane | 30 | 5052 | 0%<sup>a</sup> |
| 5512 | PR 2015 | SC | Near term | 121 | Amniotic membrane | 121 | 5318 | 0%<sup>a</sup> |
| 5803 | WT + PR 2015 | IV + IA | Death | 42 | Fetal lung | 42 | 14467 | 0%<sup>a</sup> |
| 5803 | WT+ PR 2015 | IV + IA | Death | 42 | Fetal ileum | 42 | 7137 | 0%<sup>a</sup> |
| 5803 | WT+ PR 2015 | IV + IA | Death | 42 | Fetal kidney | 42 | 12886 | 0%<sup>a</sup> |
| 5803 | WT + PR 2015 | IV + IA | Death | 42 | Fetal brain – parietal lobe | 42 | 5491 | 0.1%<sup>a</sup> |
| 5803 | WT + PR 2015 | IV + IA | Death | 42 | Placenta full thickness | 42 | 11923 | 0%<sup>a</sup> |
| 5803 | WT + PR 2015 | IV + IA | Death | 42 | Amniotic membrane | 42 | 4935 | 0%<sup>a</sup> |

**Supplemental Table 3.** Deep sequencing data showing the percentage of reads at ZIKV 1404 in non-pregnant rhesus macaques. ‘1:1 Mix’ denotes an equal mixture of ZIKV M1404 and I1404. LN is lymph node. DPI is days post-inoculation. SC is subcutaneous. These same data are shown graphically in Figure 2E.

| ANIMAL IDENTIFIER or INOCULUM | ZIKV STRAIN | INOCULATION ROUTE | TISSUE | DPI | NUCLEOTIDE COVERAGE AT 1404 | % I1404 MUTANT |
| --- | --- | --- | --- | --- | --- | --- |
| INOCULUM | 1:1 Mix | N/A | N/A | N/A | 1013 | 51% |
| 5123 | 1:1 Mix | SC | Plasma | 3 | 7920 | 18% |
| 5123 | 1:1 Mix | SC | Plasma | 5 | 7 | 14% |
| 5123 | 1:1 Mix | SC | Inguinal LN | 15 | 1607 | 32% |
| 5123 | 1:1 Mix | SC | Mesenteric LN | 15 | 881 | 22% |
| 5123 | 1:1 Mix | SC | Spleen | 15 | 170 | 25% |
| 5779 | 1:1 Mix | SC | Plasma | 3 | 7922 | 27% |
| 5779 | 1:1 Mix | SC | Plasma | 5 | 7888 | 21% |
| 5779 | 1:1 Mix | SC | Ileum | 15 | 384 | 9% |
| 5779 | 1:1 Mix | SC | Inguinal LN | 15 | 412 | 50% |
| 5779 | 1:1 Mix | SC | Mesenteric LN | 15 | 126 | 45% |
| 5779 | 1:1 Mix | SC | Spleen | 15 | 496 | 31% |

## REFERENCES

1. Duffy MR, Chen TH, Hancock WT, Powers AM, Kool JL, Lanciotti RS, Pretrick M, Marfel M, Holzbauer S, Dubray C, Guillaumot L, Griggs A, Bel M, Lambert AJ, Laven J, Kosoy O, Panella A, Biggerstaff BJ, Fischer M, Hayes EB. 2009. Zika virus outbreak on Yap Island, Federated States of Micronesia. N Engl J Med2009/06/12. 360:2536–2543.

2. Hayes EB. 2009. Zika virus outside Africa. Emerg Infect Dis 15:1347–1350.

3. Martines RB, Bhatnagar J, de Oliveira Ramos AM, Davi HPF, Iglezias SDA, Kanamura CT, Keating MK, Hale G, Silva-Flannery L, Muehlenbachs A, Ritter J, Gary J, Rollin D, Goldsmith CS, Reagan-Steiner S, Ermias Y, Suzuki T, Luz KG, de Oliveira WK, Lanciotti R, Lambert A, Shieh WJ, Zaki SR. 2016. Pathology of congenital Zika syndrome in Brazil: a case series. Lancet 388:898–904.

4. de Noronha L, Zanluca C, Azevedo MLV, Luz KG, dos Santos CND. 2016. Zika virus damages the human placental barrier and presents marked fetal neurotropism. Mem Inst Oswaldo Cruz 111:287–293.

5. Brasil P, Pereira Jr. JP, Raja Gabaglia C, Damasceno L, Wakimoto M, Ribeiro Nogueira RM, Carvalho de Sequeira P, Machado Siqueira A, Abreu de Carvalho LM, Cotrim da Cunha D, Calvet GA, Neves ES, Moreira ME, Rodrigues Baiao AE, Nassar de Carvalho PR, Janzen C, Valderramos SG, Cherry JD, Bispo de Filippis AM, Nielsen-Saines K. 2016. Zika Virus Infection in Pregnant Women in Rio de Janeiro - Preliminary Report. N Engl J Med.

6. Oliveira Melo AS, Malinger G, Ximenes R, Szejnfeld PO, Alves Sampaio S, Bispo de Filippis AM. 2016. Zika virus intrauterine infection causes fetal brain abnormality and microcephaly: tip of the iceberg? Ultrasound Obs Gynecol 47:6–7.

7. Costello A, Dua T, Duran P, Gulmezoglu M, Oladapo OT, Perea W, Pires J, Ramon-Pardo P, Rollins N, Saxena S. 2016. Defining the syndrome associated with congenital Zika virus infection. Bull World Heal Organ 94:406–406A.

8. Soares de Oliveira-Szejnfeld P, Levine D, Melo AS de O, Amorim MMR, Batista AGM, Chimelli L, Tanuri A, Aguiar RS, Malinger G, Ximenes R, Robertson R, Szejnfeld J, Tovar-Moll F. 2016. Congenital Brain Abnormalities and Zika Virus: What the Radiologist Can Expect to See Prenatally and Postnatally. Radiology 281:203–218.

9. Chimelli L, Melo ASO, Avvad-Portari E, Wiley CA, Camacho AHS, Lopes VS, Machado HN, Andrade C V, Dock DCA, Moreira ME, Tovar-Moll F, Oliveira-Szejnfeld PS, Carvalho ACG, Ugarte ON, Batista AGM, Amorim MMR, Melo FO, Ferreira TA, Marinho JRL, Azevedo GS, Leal J, da Costa RFM, Rehen S, Arruda MB, Brindeiro RM, Delvechio R, Aguiar RS, Tanuri A. 2017. The spectrum of neuropathological changes associated with congenital Zika virus infection. Acta Neuropathol 133:983–999.

10. França GVA, Schuler-Faccini L, Oliveira WK, Henriques CMP, Carmo EH, Pedi VD, Nunes ML, Castro MC, Serruya S, Silveira MF, Barros FC, Victora CG. 2016. Congenital Zika virus syndrome in Brazil: a case series of the first 1501 livebirths with complete investigation. Lancet 388:891–897.

11. Krauer F, Riesen M, Reveiz L, Oladapo OT, Martínez-Vega R, Porgo T V., Haefliger A, Broutet NJ, Low N. 2017. Zika Virus Infection as a Cause of Congenital Brain Abnormalities and Guillain–Barré Syndrome: Systematic Review. PLoS Med 14:1–27.

12. Sarno M, Sacramento GA, Khouri R, do Rosario MS, Costa F, Archanjo G, Santos LA, Nery Jr. N, Vasilakis N, Ko AI, de Almeida AR. 2016. Zika Virus Infection and Stillbirths: A Case of Hydrops Fetalis, Hydranencephaly and Fetal Demise. PLoS Negl Trop Dis 10:e0004517.

13. Mlakar J, Korva M, Tul N, Popovic M, Poljsak-Prijatelj M, Mraz J, Kolenc M, Resman Rus K, Vesnaver Vipotnik T, Fabjan Vodusek V, Vizjak A, Pizem J, Petrovec M, Avsic Zupanc T. 2016. Zika Virus Associated with Microcephaly. N Engl J Med 374:951–958.

14. Driggers RWW, Ho C-YY, Korhonen EMM, Kuivanen S, Jaaskelainen AJ, Smura T, Rosenberg A, Hill DAA, DeBiasi RLL, Vezina G, Timofeev J, Rodriguez FJJ, Levanov L, Razak J, Iyengar P, Hennenfent A, Kennedy R, Lanciotti R, du Plessis A, Vapalahti O, Jääskeläinen AJ, Smura T, Rosenberg A, Hill DAA, DeBiasi RLL, Vezina G, Timofeev J, Rodriguez FJJ, Levanov L, Razak J, Iyengar P, Hennenfent A, Kennedy R, Lanciotti R, du Plessis A, Vapalahti O. 2016. Zika Virus Infection with Prolonged Maternal Viremia and Fetal Brain Abnormalities. N Engl J Med 374:2142–2151.

15. Metsky HC, Matranga CB, Wohl S, Schaffner SF, Freije CA, Winnicki SM, West K, Qu J, Baniecki ML, Gladden-Young A, Lin AE, Tomkins-Tinch CH, Ye SH, Park DJ, Luo CY, Barnes KG, Shah RR, Chak B, Barbosa-Lima G, Delatorre E, Vieira YR, Paul LM, Tan AL, Barcellona CM, Porcelli MC, Vasquez C, Cannons AC, Cone MR, Hogan KN, Kopp EW, Anzinger JJ, Garcia KF, Parham LA, Ramirez RMG, Montoya MCM, Rojas DP, Brown CM, Hennigan S, Sabina B, Scotland S, Gangavarapu K, Grubaugh ND, Oliveira G, Robles-Sikisaka R, Rambaut A, Gehrke L, Smole S, Halloran ME, Villar L, Mattar S, Lorenzana I, Cerbino-Neto J, Valim C, Degrave W, Bozza PT, Gnirke A, Andersen KG, Isern S, Michael SF, Bozza FA, Souza TML, Bosch I, Yozwiak NL, MacInnis BL, Sabeti PC. 2017. Zika virus evolution and spread in the Americas. Nature 546:411–415.

16. Tsetsarkin KA, Kenney H, Chen R, Liu G, Manukyan H, Whitehead SS, Laassri M, Chumakov K, Pletnev AG. 2016. A Full-Length Infectious cDNA Clone of Zika Virus from the 2015 Epidemic in Brazil as a Genetic Platform for Studies of Virus-Host Interactions and Vaccine Development. MBio 7.

17. Yuan L, Huang XY, Liu ZY, Zhang F, Zhu XL, Yu JY, Ji X, Xu YP, Li G, Li C, Wang HJ, Deng YQ, Wu M, Cheng ML, Ye Q, Xie DY, Li XF, Wang X, Shi W, Hu B, Shi PY, Xu Z, Qin CF. 2017. A single mutation in the prM protein of Zika virus contributes to fetal microcephaly. Science 358(6365):933–936.

18. Jaeger AS, Murrieta RA, Goren LR, Crooks CM, Moriarty V, Weiler AM, Rybarczyk S, Semler MR, Id CH, Mejia A, Simmons HA, Id MF, Osorio E, Eickhoff JC, Connor SLO, Ebel GD, Friedrich C, Id MTA. 2019. Zika viruses of African and Asian lineages cause fetal harm in a mouse model of vertical transmission PLoS Negl Trop Dis : 13(4):e0007343. 1–18.

19. Shan C, Xia H, Haller SL, Azar SR, Liu Y, Liu J, Muruato AE, Chen R, Rossi SL, Wakamiya M, Vasilakis N, Pei R, Fontes-Garfias CR, Singh SK, Xie X, Weaver SC, Shi P-Y. 2020. A Zika virus envelope mutation preceding the 2015 epidemic enhances virulence and fitness for transmission. 2020. Proc Natl Acad Sci USA doi/10.1073/pnas.2005722117

20. Meaney-Delman D, Oduyebo T, Polen KN, White JL, Bingham AM, Slavinski SA, Heberlein-Larson L, St George K, Rakeman JL, Hills S, Olson CK, Adamski A, Culver Barlow L, Lee EH, Likos AM, Munoz JL, Petersen EE, Dufort EM, Dean AB, Cortese MM, Santiago GA, Bhatnagar J, Powers AM, Zaki S, Petersen LR, Jamieson DJ, Honein MA, Group USZPRPVW. 2016. Prolonged Detection of Zika Virus RNA in Pregnant Women. Obs Gynecol 128:724–730.

21. Akolekar R, Beta J, Picciarelli G, Ogilvie C, Antonio FD. Procedure-related risk of miscarriage following amniocentesis and chorionic villus sampling?: a systematic review and meta-analysis. Ultrasound in Obstetrics and Gynecology (2015) 45(1) 16–26.

22. Nguyen SM, Antony KM, Dudley DM, Kohn S, Simmons HA, Wolfe B, Salamat MS, Teixeira LBC, Wiepz GJ, Thoong TH, Aliota MT, Weiler AM, Barry GL, Weisgrau KL, Vosler LJ, Mohns MS, Breitbach ME, Stewart LM, Rasheed MN, Newman CM, Graham ME, Wieben OE, Turski PA, Johnson KM, Post J, Hayes JM, Schultz-Darken N, Schotzko ML, Eudailey JA, Permar SR, Rakasz EG, Mohr EL, Capuano 3rd S, Tarantal AF, Osorio JE, O’Connor SL, Friedrich TC, O’Connor DH, Golos TG. 2017. Highly efficient maternal-fetal Zika virus transmission in pregnant rhesus macaques. PLoS Pathog 13:e1006378.

23. Coffey LL, Pesavento PA, Keesler RI, Singapuri A, Watanabe J, Watanabe R, et al. Zika Virus Tissue and Blood Compartmentalization in Acute Infection of Rhesus Macaques. PLoS One. 2017;12: e0171148.

24. Coffey LL, Keesler RI, Pesavento PP, Woolard K, Singapuri A, Watanabe J, Cruzen C, Christe KL, Usachenko JU, Yee J, Heng VA, Bliss-Moreau E, Reader JR, von Morgenland W, Gibbons AM, Jackson K, Ardeshir A, Heimsath H, Permar SR, Senthamaraikannan P, Presicce P, Kallapur SG, Linnen JM, Gao K, Orr R, MacGill T, McClure M, McFarland R, Morrison JM, Van Rompay KKA. 2018. Intraamniotic Zika Virus Inoculation of Pregnant Rhesus Macaques Produces Fetal Neurologic Disease. Nat Commun (9) 2414

25. Dudley DM, van Rompay KK, Coffey LL, Ardeshir A, Keesler RI, Bliss-Moreau E, Grigsby PL, Steinbach RJ, Hirsch AJ, MacAllister RP, Pecoraro HL, Colgin LM, Hodge T, Streblow DN, Tardif S, Patterson JL, Tamhankar M, Seferovic M, Aagaard KM, Martín CSS, Chiu CY, Panganiban AT, Veazey RS, Wang X, Maness NJ, Gilbert MH, Bohm RP, Adams Waldorf KM, Gale M, Rajagopal L, Hotchkiss CE, Mohr EL, Capuano S V., Simmons HA, Mejia A, Friedrich TC, Golos TG, O’Connor DH. 2018. Miscarriage and stillbirth following maternal Zika virus infection in nonhuman primates. Nat Med. 24:1104–1107.

26. Dudley DM, Aliota MT, Mohr EL, Weiler AM, Lehrer-Brey G, Weisgrau KL, Mohns MS, Breitbach ME, Rasheed MN, Newman CM, Gellerup DD, Moncla LH, Post J, Schultz-Darken N, Schotzko ML, Hayes JM, Eudailey JA, Moody MA, Permar SR, O’Connor SL, Rakasz EG, Simmons HA, Capuano S, Golos TG, Osorio JE, Friedrich TC, O’Connor DH. 2016. A rhesus macaque model of Asian-lineage Zika virus infection. Nat Commun 7:12204.

27. Mohr EL, Block LN, Newman CM, Stewart LM, Koenig M, Semler M, Breitbach ME, Teixeira LBC, Zeng X, Weiler AM, Barry GL, Thoong TH, Wiepz GJ, Dudley DM, Simmons HA, Mejia A, Morgan TK, Salamat MS, Kohn S, Antony KM, Aliota MT, Mohns MS, Hayes JM, Schultz-Darken N, Schotzko ML, Peterson E, Capuano 3rd S, Osorio JE, O’Connor SL, Friedrich TC, O’Connor DH, Golos TG. 2018. Ocular and uteroplacental pathology in a macaque pregnancy with congenital Zika virus infection. PLoS One 13:e0190617.

28. Koide F, Goebel S, Snyder B, Walters KB, Gast A, Hagelin K, Kalkeri R, Rayner J. 2016. Development of a zika virus infection model in cynomolgus macaques. Front Microbiol 7:1–8.

29. Osuna CE, Lim SY, Deleage C, Griffin BD, Stein D, Schroeder LT, Omange R, Best K, Luo M, Hraber PT, Andersen-Elyard H, Ojeda EF, Huang S, Vanlandingham DL, Higgs S, Perelson AS, Estes JD, Safronetz D, Lewis MG, Whitney JB. 2016. Zika viral dynamics and shedding in rhesus and cynomolgus macaques. Nat Med. (12):1448–1455.

30. Waldorf KMA, Nelson BR, Stencel-Baerenwald JE, Studholme C, Kapur RP, Armistead B, Walker CL, Merillat S, Vornhagen J, Tisoncik-Go J, Baldessari A, Coleman M, Dighe MK, Shaw DWWW, Roby JA, Santana-Ufret V, Boldenow E, Li J, Gao X, Davis MA, Swanstrom JA, Jensen K, Widman DG, Baric RS, Medwid JT, Hanley KA, Ogle J, Gough GM, Lee W, English C, Durning WMI, Thiel J, Gatenby C, Dewey EC, Fairgrieve MR, Hodge RD, Grant RF, Kuller L, Dobyns WB, Hevner RF, Gale M J, Rajagopal L, Adams Waldorf KM, Nelson BR, Stencel-Baerenwald JE, Studholme C, Kapur RP, Armistead B, Walker CL, Merillat S, Vornhagen J, Tisoncik-Go J, Baldessari A, Coleman M, Dighe MK, Shaw DWWW, Roby JA, Santana-Ufret V, Boldenow E, Li J, Gao X, Davis MA, Swanstrom JA, Jensen K, Widman DG, Baric RS, Medwid JT, Hanley KA, Ogle J, Gough GM, Lee W, English C, Durning WMI, Thiel J, Gatenby C, Dewey EC, Fairgrieve MR, Hodge RD, Grant RF, Kuller L, Dobyns WB, Hevner RF, Gale M, Rajagopal L. 2018. Congenital Zika virus infection as a silent pathology with loss of neurogenic output in the fetal brain. Nat Med. 2016;22: 1256–1259.

31. Adams Waldorf KM, Stencel-Baerenwald JE, Kapur RPR, Studholme C, Boldenow E, Vornhagen J, Baldessari A, Dighe MK, Thiel J, Merillat S, Armistead B, Tisoncik-Go J, Green RGR, Davis MA, Dewey EC, Fairgrieve MR, Gatenby JC, Richards T, Garden GA, Diamond MSM, Juul SES, Grant RF, Kuller L, Shaw DWWW, Ogle J, Gough GM, Lee W, English C, Hevner RF, Dobyns WB, Gale Jr M, Rajagopal L, Gale M, Rajagopal L. 2016. Fetal brain lesions after subcutaneous inoculation of Zika virus in a pregnant nonhuman primate. Nat Med 22:1256–1259.

32. Aid M, Abbink P, Larocca RA, Boyd M, Nityanandam R, Nanayakkara O, Martinot AJ, Moseley ET, Blass E, Borducchi EN, Chandrashekar A, Brinkman AL, Molloy K, Jetton D, Tartaglia LJ, Liu J, Best K, Perelson AS, De La Barrera RA, Lewis MG, Barouch DH. 2017. Zika Virus Persistence in the Central Nervous System and Lymph Nodes of Rhesus Monkeys. Cell 169:610-620.e14.

33. Hirsch AJ, Smith JL, Haese NN, Broeckel RM, Parkins CJ, Kreklywich C, DeFilippis VR, Denton M, Smith PP, Messer WB, Colgin LMA, Ducore RM, Grigsby PL, Hennebold JD, Swanson T, Legasse AW, Axthelm MK, MacAllister R, Wiley CA, Nelson JA, Streblow DN. 2017. Zika Virus infection of rhesus macaques leads to viral persistence in multiple tissues. PLoS Pathog 13:1–23.

34. Van Rompay KKA, Keesler RI, Ardeshir A, Watanabe J, Usachenko J, Singapuri A, Cruzen C, Bliss-Moreau E, Murphy AM, Yee JL, Webster H, Dennis M, Singh T, Heimsath H, Lemos D, Stuart J, Morabito KM, Foreman BM, Burgomaster KE, Noe AT, Dowd KA, Ball E, Woolard K, Presicce P, Kallapur SG, Permar SR, Foulds KE, Coffey LL, Pierson TC, Graham BS. 2019. DNA vaccination before conception protects Zika virus-exposed pregnant macaques against prolonged viremia and improves fetal outcomes. Sci Transl Med 11:1–12.

35. Van Rompay KKA, Keesler RI, Ardeshir A, Watanabe J, Usachenko J, Singapuri A, Cruzen C, Bliss-Moreau E, Murphy AM, Yee JAL, Webster H, Dennis M, Singh T, Heimsath H, Lemos D, Stuart J, Morabito KM, Foreman BM, Burgomaster KE, Noe AT, Dowd KA, Ball E, Woolard K, Presicce P, Kallapur SG, Permar SR, Foulds KE, Coffey LL, Pierson TC, Graham BS. 2019. DNA vaccination before conception protects Zika virus– exposed pregnant macaques against prolonged viremia and improves fetal outcomes. Sci Transl Med. doi:10.1126/scitranslmed.aay2736

36. Brown RJ, Peters PJ, Caron C, Gonzalez-Perez MP, Stones L, Ankghuambom C, Pondei K, McClure CP, Alemnji G, Taylor S, Sharp PM, Clapham PR, Ball JK. 2011. Intercompartmental recombination of HIV-1 contributes to env intrahost diversity and modulates viral tropism and sensitivity to entry inhibitors. J Virol 85:6024–6037.

37. Frost SD, Wrin T, Smith DM, Kosakovsky Pond SL, Liu Y, Paxinos E, Chappey C, Galovich J, Beauchaine J, Petropoulos CJ, Little SJ, Richman DD. 2005. Neutralizing antibody responses drive the evolution of human immunodeficiency virus type 1 envelope during recent HIV infection. Proc Natl Acad Sci U S A 102:18514–18519.

38. Leslie AJ, Pfafferott KJ, Chetty P, Draenert R, Addo MM, Feeney M, Tang Y, Holmes EC, Allen T, Prado JG, Altfeld M, Brander C, Dixon C, Ramduth D, Jeena P, Thomas SA, St John A, Roach TA, Kupfer B, Luzzi G, Edwards A, Taylor G, Lyall H, Tudor-Williams G, Novelli V, Martinez-Picado J, Kiepiela P, Walker BD, Goulder PJ. 2004. HIV evolution: CTL escape mutation and reversion after transmission. Nat Med 10:282–289.

39. Vignuzzi M, Stone JK, Arnold JJ, Cameron CE, Andino R. 2006. Quasispecies diversity determines pathogenesis through cooperative interactions in a viral population. Nature2005/12/06. 439:344–348.

40. Borucki MK, Allen JE, Chen-Harris H, Zemla A, Vanier G, Mabery S, Torres C, Hullinger P, Slezak T. 2013. The role of viral population diversity in adaptation of bovine coronavirus to new host environments. PLoS One 8:e52752.

41. Nora T, Bouchonnet F, Labrosse B, Charpentier C, Mammano F, Clavel F, Hance AJ. 2008. Functional diversity of HIV-1 envelope proteins expressed by contemporaneous plasma viruses. Retrovirology 5:23.

42. Memoli MJ, Bristol T, Proudfoot KE, Davis AS, Dunham EJ, Taubenberger JK. 2012. In vivo evaluation of pathogenicity and transmissibility of influenza A(H1N1)pdm09 hemagglutinin receptor binding domain 222 intrahost variants isolated from a single immunocompromised patient. Virology 428:21–29.

43. Domingo E, Gomez J. 2007. Quasispecies and its impact on viral hepatitis. Virus Res 127:131–150.

44. Debbink K, Lindesmith LC, Donaldson EF, Swanstrom J, Baric RS. 2014. Chimeric GII.4 norovirus virus-like-particle-based vaccines induce broadly blocking immune responses. J Virol 88:7256–7266.

45. Norstrom MM, Buggert M, Tauriainen J, Hartogensis W, Prosperi MC, Wallet MA, Hecht FM, Salemi M, Karlsson AC. 2012. Combination of immune and viral factors distinguishes low-risk versus high-risk HIV-1 disease progression in HLA-B*5701 subjects. J Virol 86:9802–9816.

46. Caporale M, Di Gialleonorado L, Janowicz A, Wilkie G, Shaw A, Savini G, Van Rijn PA, Mertens P, Di Ventura M, Palmarini M. 2014. Virus and host factors affecting the clinical outcome of bluetongue virus infection. J Virol 88:10399–10411.

47. Riemersma KK, Steiner C, Singapuri A, Coffey LL. Chikungunya Virus Fidelity Variants Exhibit Differential Attenuation and Population Diversity in Cell Culture and Mice. J Virol. 2019;93: 1–19.

48. Riemersma KK, Coffey LL. 2019. Chikungunya virus populations experience diversity-dependent attenuation and purifying intra-vector selection in Californian Aedes aegypti mosquitoes. PLoS Negl Trop Dis. 2019. doi:10.1371/journal.pntd.0007853

49. Coffey LL, Beeharry Y, Borderia A V, Blanc H, Vignuzzi M. 2011. Arbovirus high fidelity variant loses fitness in mosquitoes and mice. Proc Natl Acad Sci U S A2011/09/08. 108:16038–16043.

50. Rozen-Gagnon K, Stapleford KA, Mongelli V, Blanc H, Failloux AB, Saleh MC, Vignuzzi M. 2014. Alphavirus mutator variants present host-specific defects and attenuation in Mammalian and insect models. PLoS Pathog2014/01/24. 10:e1003877.

51. Magnani DM, Rogers TF, Maness NJ, Grubaugh ND, Beutler N, Bailey VK, Gonzalez-Nieto L, Gutman MJ, Pedreño-Lopez N, Kwal JM, Ricciardi MJ, Myers TA, Julander JG, Bohm RP, Gilbert MH, Schiro F, Aye PP, Blair R V., Martins MA, Falkenstein KP, Kaur A, Curry CL, Kallas EG, Desrosiers RC, Goldschmidt-Clermont PJ, Whitehead SS, Andersen KG, Bonaldo MC, Lackner AA, Panganiban AT, Burton DR, Watkins DI. 2018. Fetal demise and failed antibody therapy during Zika virus infection of pregnant macaques. Nat Commun 9:1–8.

52. Pfeiffer F, Gröber C, Blank M, Händler K, Beyer M, Schultze JL, Mayer G. 2018. Systematic evaluation of error rates and causes in short samples in next-generation sequencing. Sci Rep. 8(1)

53. Vermillion MS, Lei J, Shabi Y, Baxter VK, Crilly NP, McLane M, Griffin DE, Pekosz A, Klein SL, Burd I. 2017. Intrauterine Zika virus infection of pregnant immunocompetent mice models transplacental transmission and adverse perinatal outcomes. Nat Commun 8:14575.

54. Weger-Lucarelli J, Rückert C, Chotiwan N, Nguyen C, Garcia Luna SM, Fauver JR, Foy BD, Perera R, Black WC, Kading RC, Ebel GD. 2016. Vector Competence of American Mosquitoes for Three Strains of Zika Virus. PLoS Negl Trop Dis 10:1–16.

55. Cao B, Diamond MS, Mysorekar IU. 2017. Maternal-fetal transmission of zika virus: Routes and signals for infection. J Interf Cytokine Res. doi:10.1089/jir.2017.0011

56. Tabata T, Petitt M, Puerta-Guardo H, Michlmayr D, Wang C, Fang-Hoover J, Harris E, Pereira L. 2016. Zika Virus Targets Different Primary Human Placental Cells, Suggesting Two Routes for Vertical Transmission. Cell Host Microbe 20:155–166.

57. Miner JJ, Diamond MS. 2017. Zika Virus Pathogenesis and Tissue Tropism. Cell Host Microbe. doi:10.1016/j.chom.2017.01.004

58. Russo FB, Jungmann P, Beltrão-Braga PCB. 2017. Zika infection and the development of neurological defects. Cell Microbiol. doi:10.1111/cmi.12744

59. CDC. 2016. Vital Signs: Preparing for Local Mosquito-Borne Transmission of Zika Virus - United States, 2016. MMWR Morb Mortal Wkly Rep 65:352.

60. Li X-D, Deng C-L, Ye H-Q, Zhang H-L, Zhang Q-Y, Chen D-D, Zhang P-T, Shi P-Y, Yuan Z-M, Zhang B. 2016. Transmembrane Domains of NS2B Contribute to both Viral RNA Replication and Particle Formation in Japanese Encephalitis Virus. J Virol. 90(12):5735–5749.

61. Li Y, Li Q, Wong YL, Liew LSY, Kang C. 2015. Membrane topology of NS2B of dengue virus revealed by NMR spectroscopy. Biochim Biophys Acta - Biomembr. (1848) 10A:2244–2252.

62. Chambers TJ, Nestorowicz A, Amberg SM, Rice CM. 1993. Mutagenesis of the yellow fever virus NS2B protein: effects on proteolytic processing, NS2B-NS3 complex formation, and viral replication. J Virol. 67(11): 6797–6807

63. Grubaugh ND, Petrone ME, Holmes EC. 2020. We shouldn’t worry when a virus mutates during disease outbreaks. Nat Microbiol. 5: 529–530.

64. Lanciotti RS, Kosoy OL, Laven JJ, Velez JO, Lambert AJ, Johnson AJ, Stanfield SM, Duffy MR. 2008. Genetic and serologic properties of Zika virus associated with an epidemic, Yap State, Micronesia, 2007. 2008. Emerg Infect Dis. 14:1232–1239.

65. Deng X, Achari A, Federman S, Yu G, Somasekar S, Bártolo I, Yagi S, Mbala-Kingebeni P, Kapetshi J, Ahuka-Mundeke S, Muyembe-Tamfum JJ, Ahmed AA, Ganesh V, Tamhankar M, Patterson JL, Ndembi N, Mbanya D, Kaptue L, McArthur C, Muñoz-Medina JE, Gonzalez-Bonilla CR, López S, Arias CF, Arevalo S, Miller S, Stone M, Busch M, Hsieh K, Messenger S, Wadford DA, Rodgers M, Cloherty G, Faria NR, Thézé J, Pybus OG, Neto Z, Morais J, Taveira N, R. Hackett J, Chiu CY. 2020. Metagenomic sequencing with spiked primer enrichment for viral diagnostics and genomic surveillance. Nat Microbiol. 5: 443–454.

66. Borucki MK, Collette NM, Coffey LL, Van Rompay KKA, Hwang MH, Thissen JB, Allen JE, Zemla AT. 2019. Multiscale analysis for patterns of Zika virus genotype emergence, spread, and consequence. PLoS One. 14(12): e0225699.

67. Grubaugh ND, Gangavarapu K, Quick J, Matteson NL, Jesus JG De, Main BJ, Tan AL, Paul LM, Brackney DE, Grewal S, Gurfield N, Rompay KKA Van, Isern S, Michael SF, Coffey LL, Loman NJ, Andersen KG. 2019. An amplicon-based sequencing framework for accurately measuring intrahost virus diversity using PrimalSeq and iVar 1–19.

68. Quick J, Grubaugh ND, Pullan ST, Claro IM, Smith AD, Gangavarapu K, Oliveira G, Robles-Sikisaka R, Rogers TF, Beutler NA, Burton DR, Lewis-Ximenez LL, De Jesus JG, Giovanetti M, Hill SC, Black A, Bedford T, Carroll MW, Nunes M, Alcantara LC, Sabino EC, Baylis SA, Faria NR, Loose M, Simpson JT, Pybus OG, Andersen KG, Loman NJ. 2017. Multiplex PCR method for MinION and Illumina sequencing of Zika and other virus genomes directly from clinical samples. Nat Protoc.

69. Li H. 2013. Aligning sequence reads, clone sequences and assembly contigs with BWA-MEM.

70. Li H. 2011. A statistical framework for SNP calling, mutation discovery, association mapping and population genetical parameter estimation from sequencing data. Bioinformatics 27:2987–2993.

